# HIV-2 evades restriction by ZAP through adaptations in the U3 LTR region despite increased CpG levels

**DOI:** 10.1101/2024.10.13.618097

**Authors:** Dorota Kmiec, Rayhane Nchioua, Alexander Gabel, Asimenia Vlachou, Sabina Ganskih, Sümeyye Erdemci-Evin, Stacey Lapp, Diane Carnathan, Steven Bosinger, Ben Berkhout, Atze T Das, Mathias Munschauer, Frank Kirchhoff

## Abstract

Simian immunodeficiency viruses infecting sooty mangabeys (SIVsmm) gave rise to nine groups of human immunodeficiency virus type 2 (HIV-2). Two of these (A and B) spread substantially with an estimated 1- 2 million individuals affected. The evolutionary adaptations that facilitated HIV-2’s spread in humans are still poorly understood. Here, we report that diverse SIVsmm strains efficiently infect primary human T cells. However, they are more sensitive to interferon than HIV-2, indicating that interferon-stimulated genes (ISGs) pose a barrier to the successful spread of SIVsmm in humans. One of the best-known antiviral ISGs is the zinc finger antiviral protein (ZAP), which targets CpG dinucleotides in RNA. To evade ZAP-mediated restriction, many viruses, including HIV-1, suppress their CpG content. Unexpectedly, we found that HIV-2 is more resistant to ZAP restriction than HIV-1 and SIVsmm despite having 33% more CpGs. Identification of ZAP binding sites using RNA eCLIP and analyses of chimeric HIV-2/SIVsmm viruses, revealed that the determinants of ZAP resistance map to the U3 region of the long-terminal repeat and promote HIV-2 replication in primary human T cells. Our results indicate that ZAP poses a barrier to SIVsmm infection in humans and that HIV-2 evolved a CpG-independent mechanism to evade it.

## INTRODUCTION

Zinc-finger antiviral protein (ZAP), also known as ARTD13, PARP13, and ZC3HAV1, is a broadly expressed cellular RNA-binding protein that inhibits multiple RNA and DNA viruses (1–3). Like other antiviral restriction factors that constitute intrinsic immunity, ZAP evolved under positive selection and its expression is upregulated by interferons (IFNs) (3–6). ZAP binds to CpG dinucleotides in HIV-1 RNAs and recruits the co-factors ubiquitin ligase Tripartite Motif Containing 25 (TRIM25) and endoribonuclease KH and NYN Domain Containing (KHNYN) to degrade it, which in turn reduces viral protein expression and replication (7–10). CpGs are the least abundant dinucleotide type found in the transcriptomes of most vertebrates, including humans (2, 11, 12). ZAP exploits this CpG suppression to distinguish between self and non-self RNAs and target viral transcripts.

The continuous arms race between viruses and their hosts shapes pathogen evolution and enables the evasion of antiviral restriction factors such as ZAP. Many successful viruses mimic the CpG suppression of their host’s genomes to evade the recognition by ZAP. For example, HIV-1 suppresses its genomic CpG content by over 80%, which is even lower than CpG suppression of the human transcriptome (60%) (7, 13). However, the CpG content of primate lentiviral genomes varies greatly and does not correlate with ZAP sensitivity (13), suggesting alternative ZAP evasion mechanisms.

Simian immunodeficiency virus (SIV) species naturally infect over 40 African primate species. Of those, SIVs infecting chimpanzees, gorillas and sooty mangabeys successfully crossed the interspecies barrier and gave rise to HIV-1 (SIVcpz, SIVgor) and HIV-2 (SIVsmm). While the HIV-1 group M originating from SIVcpz has spread globally causing >95% of all HIV infections, HIV-2 infects around 1-2 million people, mostly in West Africa (14). HIV-1 and HIV-2 evolved sophisticated ways to evade or counteract human immune responses and cause persistent infection that, if untreated, eventually leads to a fatal acquired immune deficiency syndrome (AIDS). SIVsmm, the ancestor of HIV-2, does not cause disease in its natural host and is highly prevalent in sooty mangabeys, which are hunted for bushmeat or kept as pets in parts of West Africa (15). SIVsmm strains readily infect human T cells *in vitro* (16–18). Therefore, it is not surprising that SIVsmm crossed the species barrier from monkeys to humans on at least nine occasions. However, only two of these zoonotic transmissions spread among humans, giving rise to epidemic HIV-2 groups A and B. While the overall prevalence of HIV-2 is declining, new infections with HIV-2 groups A, B, and their recombined form (AB) continue to be reported outside of West Africa (19–21). It remains to be defined what type of evolutionary changes the epidemic groups of HIV-2 gained during their human adaptation.

Here, we examined whether ZAP poses a barrier to effective spread of SIVsmm in human cells. We found that SIVsmm strains are more sensitive to IFN than HIV-2 and also significantly more susceptible to inhibition by ZAP in human T cells. Surprisingly, epidemic HIV-2 evolved resistance to ZAP despite acquiring CpG dinucleotides during its adaptation to humans. Through enhanced crosslinking immunoprecipitation (eCLIP) analysis of ZAP-binding sites in viral RNA and exchanging specific genomic regions between HIV-2 and SIVsmm, we show that the factors responsible for HIV-2’s evasion of ZAP do not map to the *env* gene coding region, as observed in HIV-1. Instead, they involve changes in the U3 region of the long terminal repeat (LTR), which enhance HIV-2 fitness in primary human T lymphocytes. Our results suggest that CpG-independent changes in the U3 region of the LTR contributed to the evolution of ZAP resistance and epidemic spread of HIV-2 in humans.

## METHODS

### Cell lines

HEK293T and HeLa cells were obtained from ATCC. HEK293T ZAP KO cells have been previously described (7). THP-1 as well as Jurkat WT and ZAP KO cells were described before (22, 23). TZM-bl cells express CD4, CCR5, and CXCR4 and contain the β-galactosidase genes under the control of the HIV promoter (24, 25).

Adherent cell lines were cultured in Dulbecco’s modified Eagle’s medium (DMEM) supplemented with 2.5% (during virus production) or 10% (at all other times) heat-inactivated foetal calf serum (FCS), 2 mM L-glutamine, 100 U/ml penicillin, and 100 μg/ml streptomycin. Jurkat cells were cultured in RPMI supplemented with 10% FCS, 2 mM L-glutamine, 100 U/ml penicillin, and 100 μg/ml streptomycin.

### Isolation of primary cells

Peripheral blood mononuclear cells (PBMCs) or CD4+ T cells from healthy human donors were isolated using lymphocyte separation medium (Biocoll separating solution; Biochrom) or RosetteSep Human CD4+ T Cell Enrichment Cocktail (Stemcell), stimulated for 3 days with phytohemagglutinin (PHA; 2 μg/ml), and 10 ng/ml interleukin-2 (IL-2) in RPMI 1640 medium containing 10% fetal calf serum. Where indicated, cells were stimulated with IFNα 24h prior to infection.

### Expression constructs

The pCG IRES eBFP vector encoding human ZAP-L containing an N-terminal hemagglutinin (HA) tag and empty vector control were described before (13). The infectious molecular clones of HIV-1 NL4-3, HIV-1 NL4-3 CpG high mutant, CH77TF, CH58 TF, THRO, CH167, HIV-2 7312A, ST, KR, GH123, ROD10, MCR, MCN, as well as SIVsmm PGM, SIVsmm PGM nef-IRES eGFP, L1.V1, L1.V2, L2, L3, L4 and L5 were described previously (8, 26–39). HA-ZAP-L zinc-finger 1-4 deletion mutant was generated by Q5 site-directed mutagenesis using pCG HA-ZAP-L IRES BFP as a template. The pCG mCherry-ZAP-L IRES BFP expression construct was generated by Gibson assembly using the XbaI/Mlul cloning sites. SIVsmm Env and Nef expression vectors and HIV-2 7312A *env* chimeras were previously reported (16). HIV-2 7312A IRES eGFP constructs carrying premature stop codons in *vpr* and *nef* ORFs were described before (31). SIVsmm Vpr and Nef expression constructs were described before (18, 40). Chimeric HIV-2 and SIVsmm infectious molecular clones as well as CpG mutants thereof were generated using Gibson assembly (NEB) and cloned into pCR XL TOPO or pBR vector using NotI/MluI restriction sites. All constructs were cloned using primers (from biomers.net) listed in Table 1 and their correctness was confirmed by Sanger sequencing (eurofins).

**Table 1.**
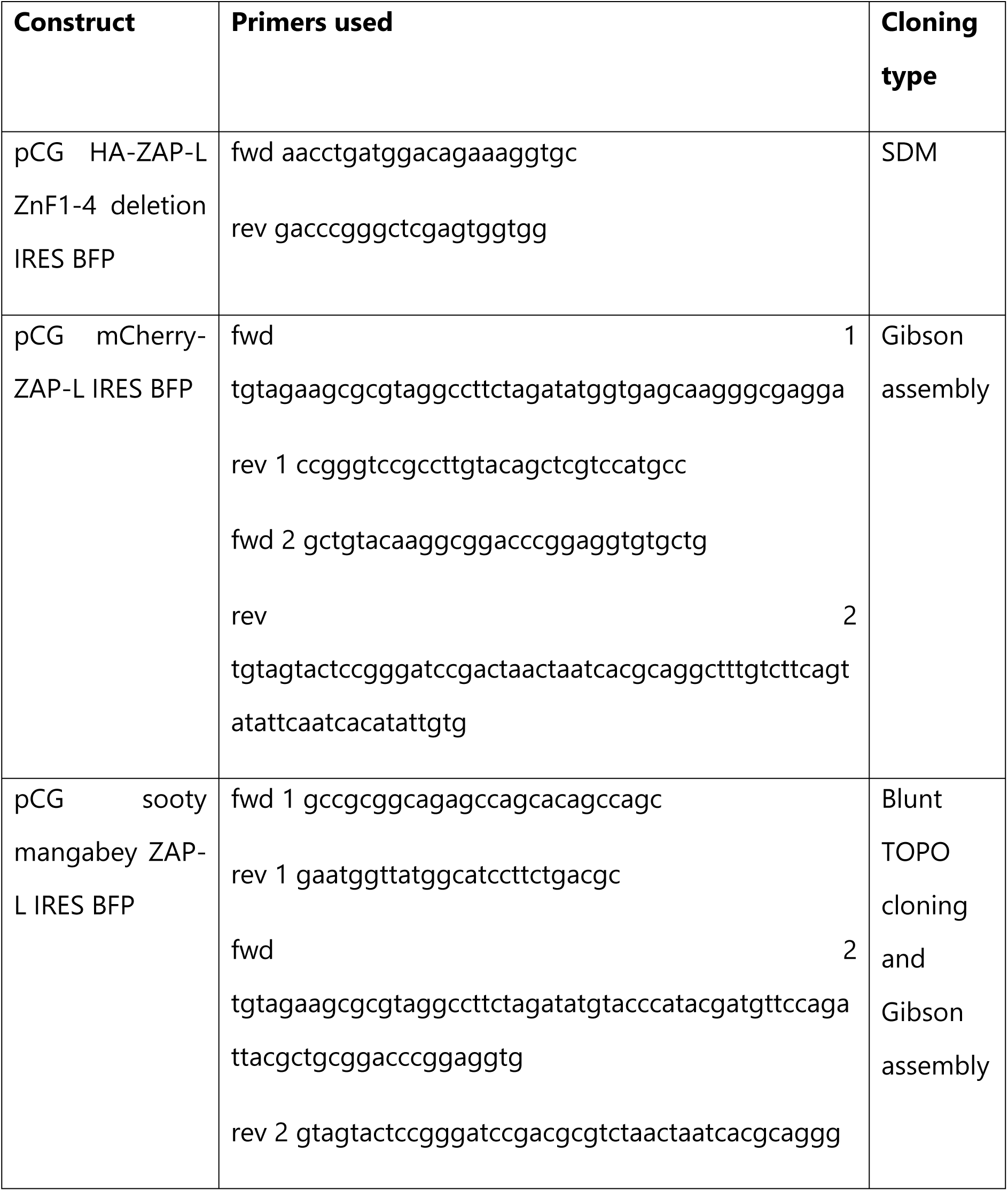
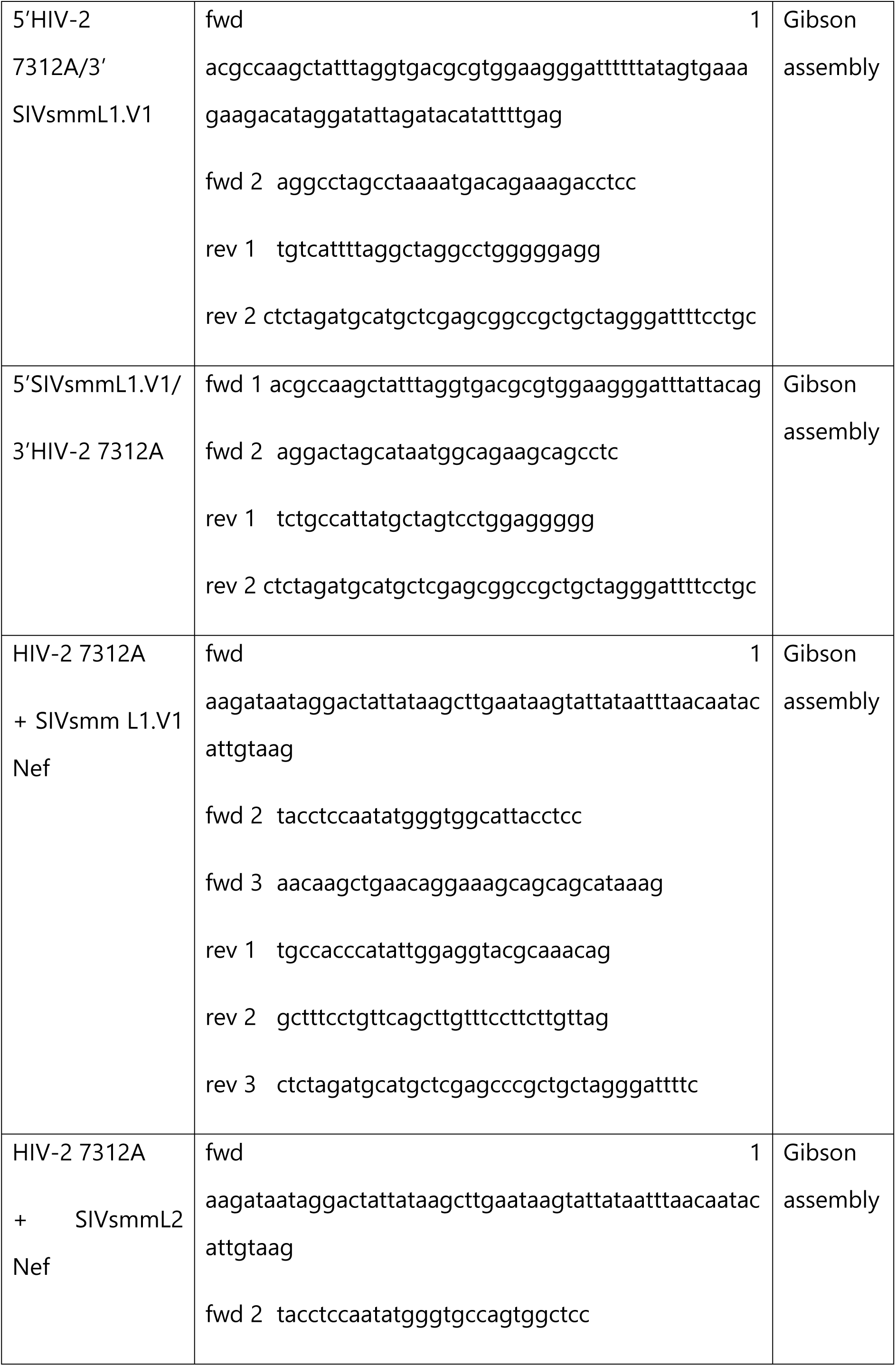
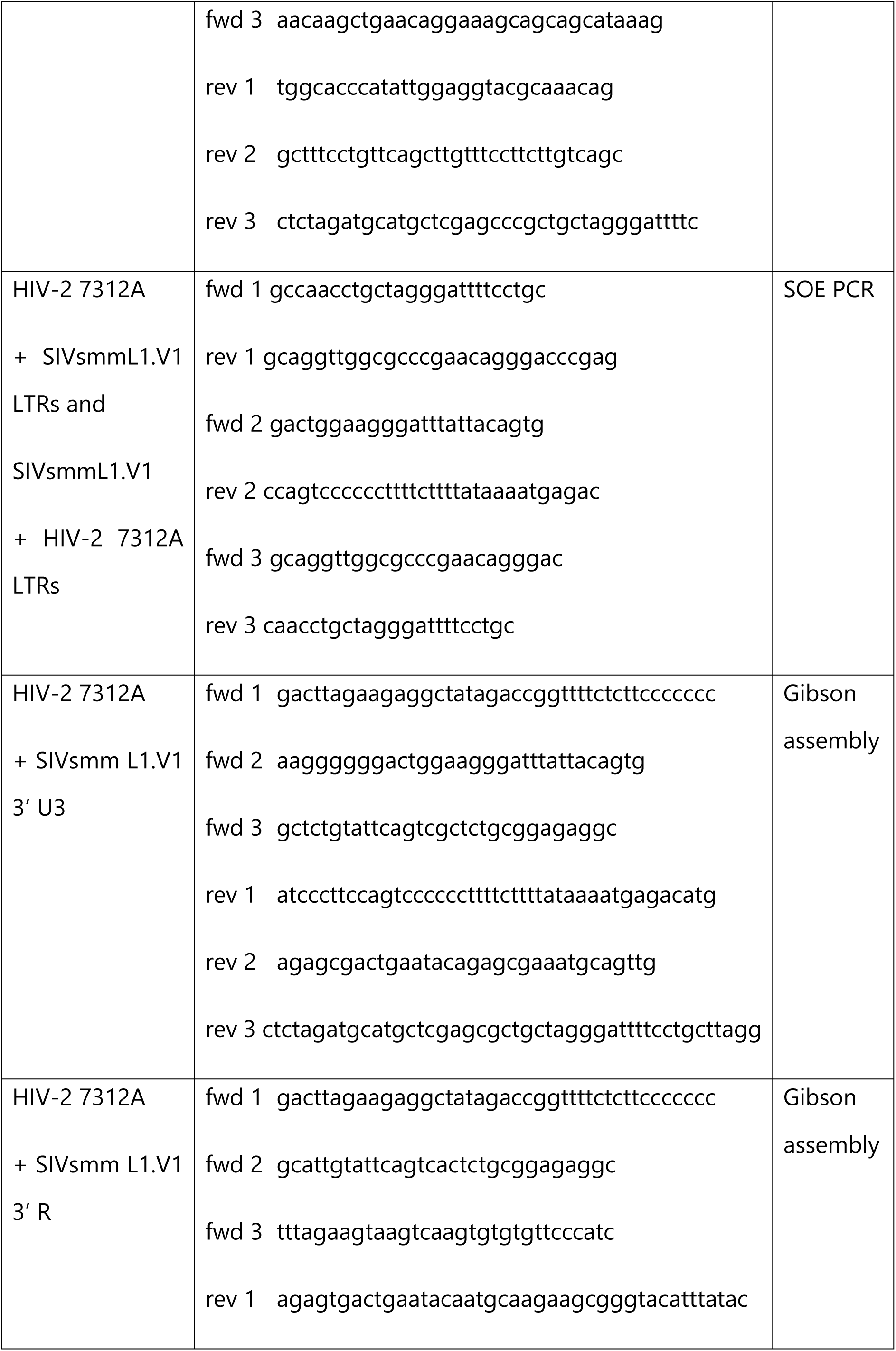
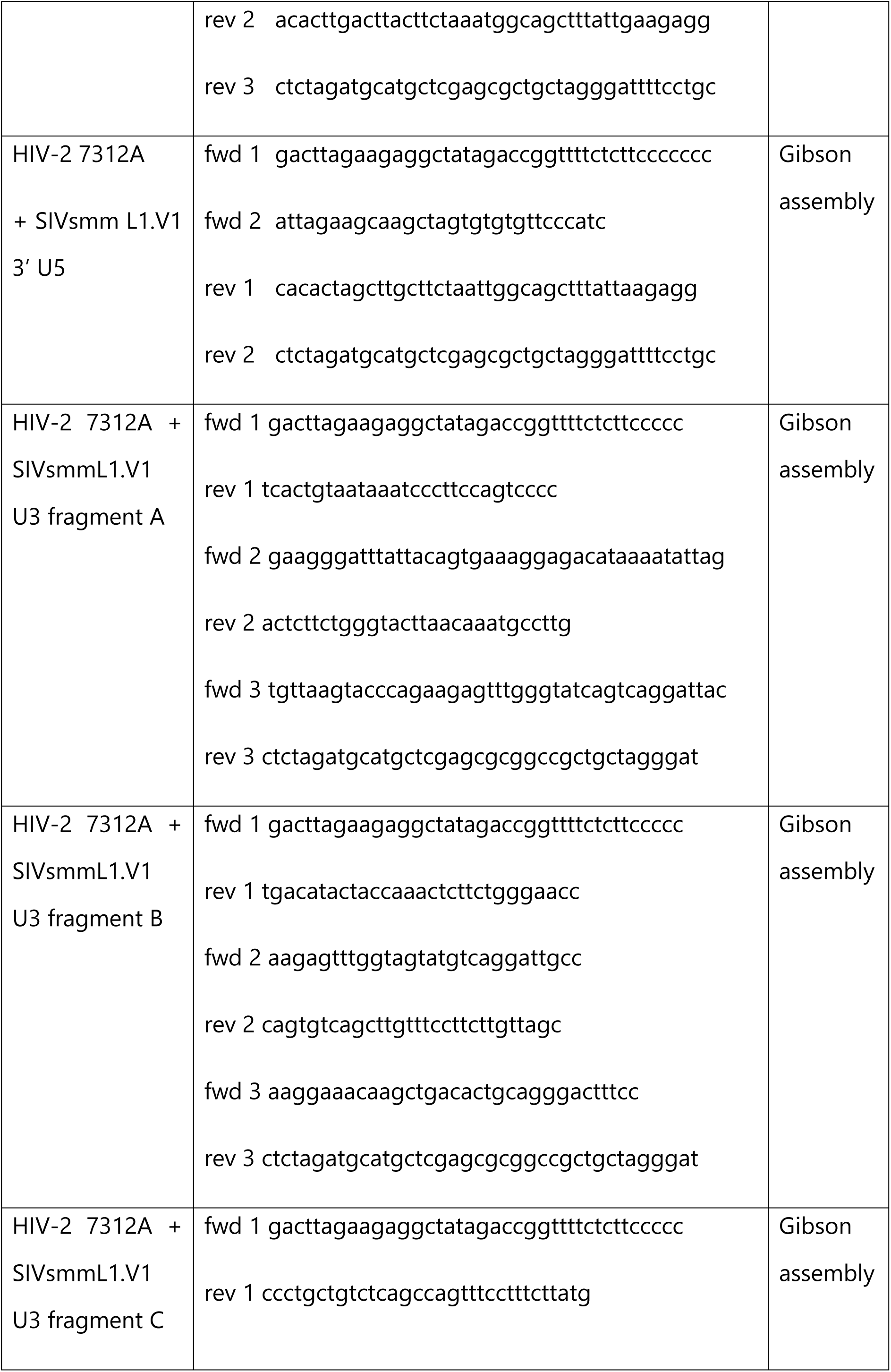
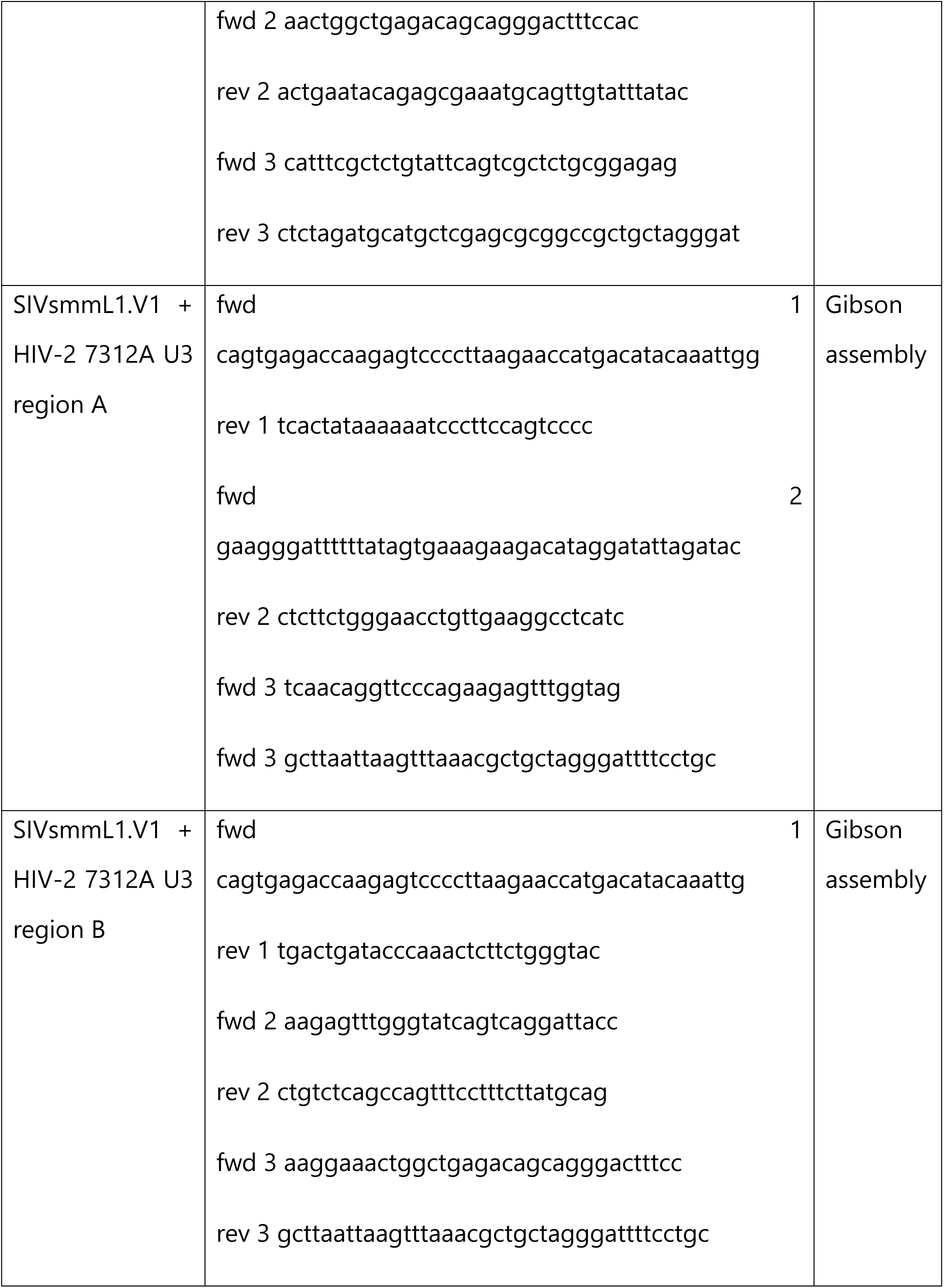
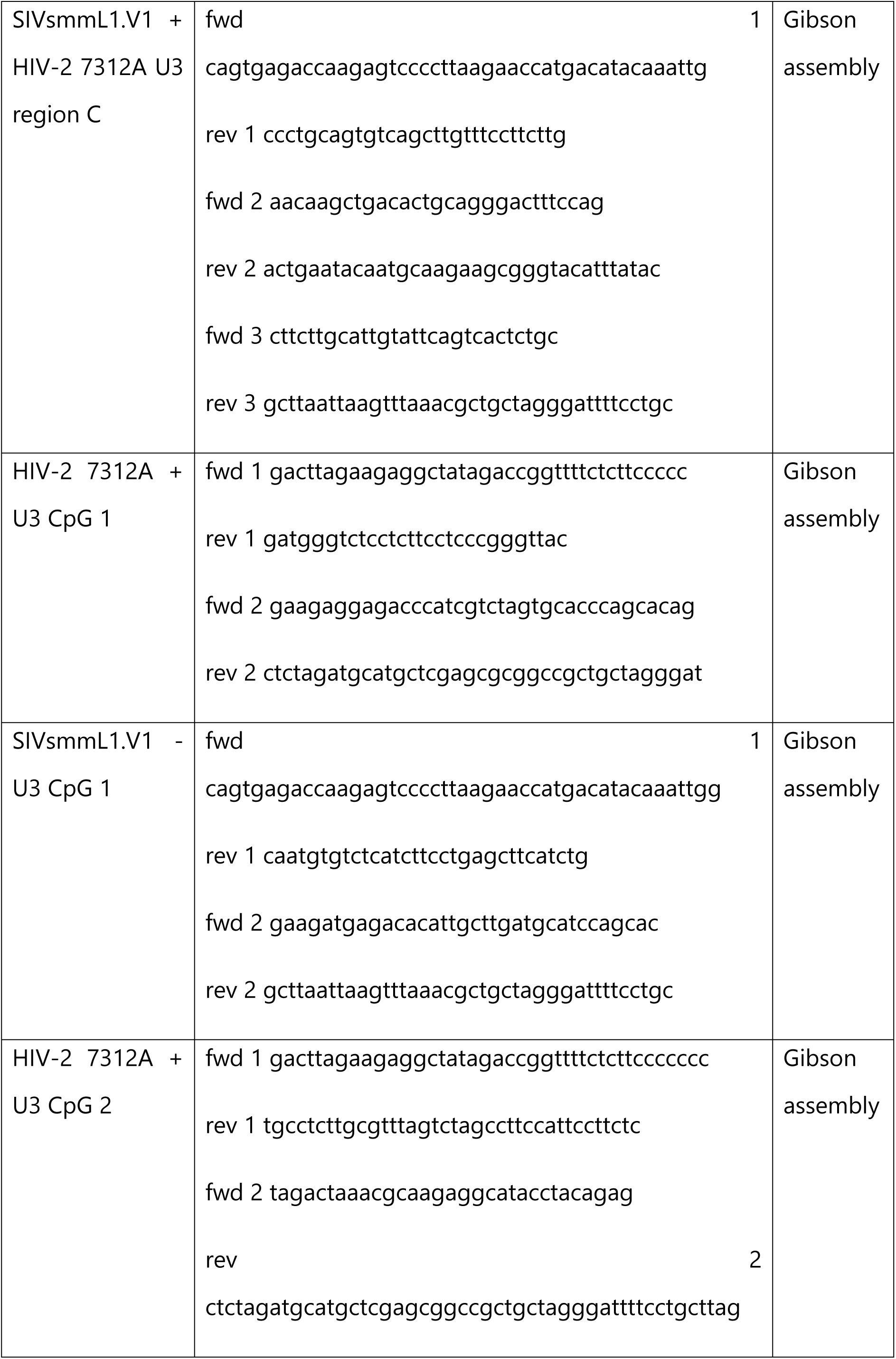
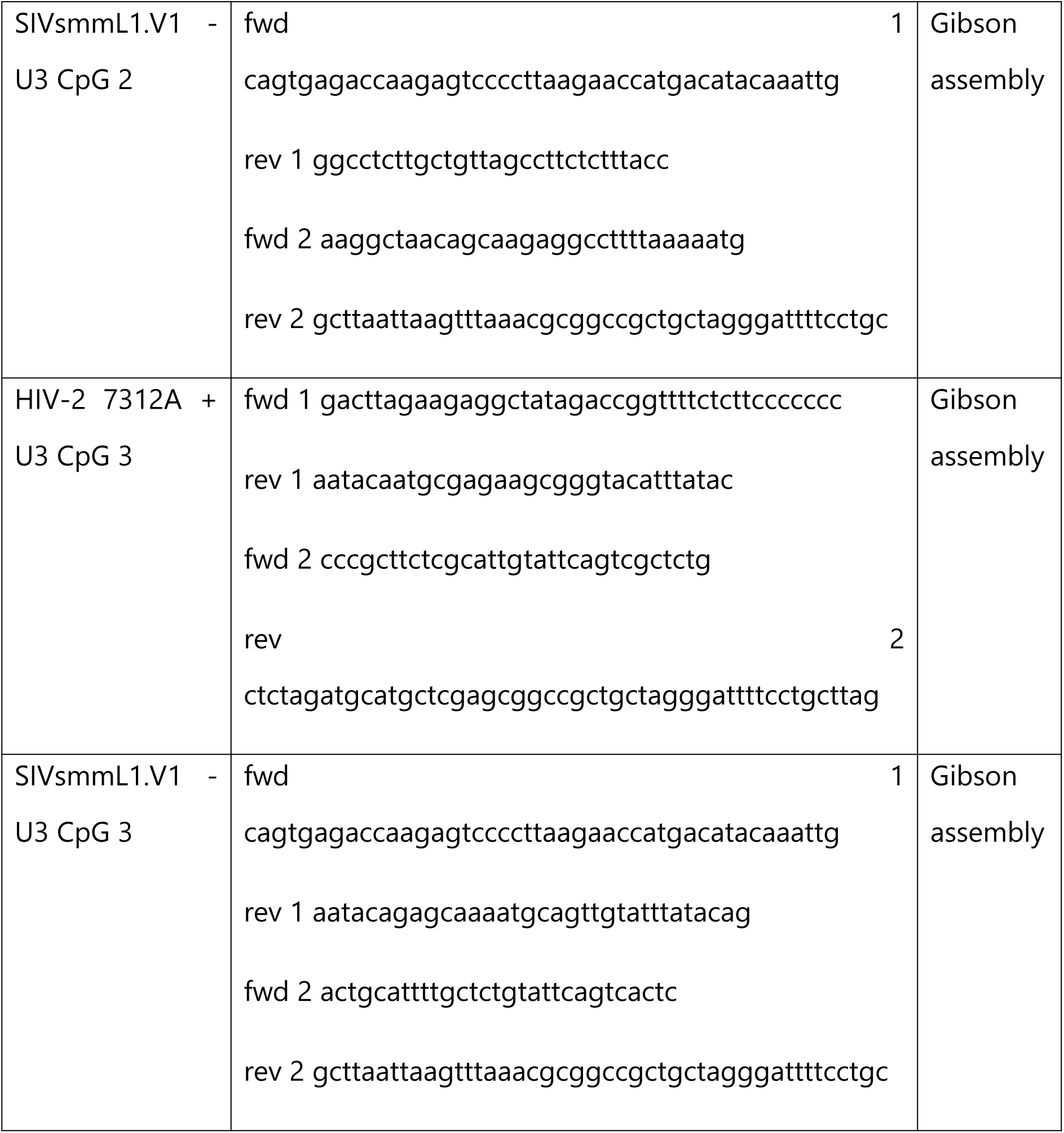
Oligonucleotides used for cloning.

### Generation of sooty mangabey ZAP

Whole blood from two sooty mangabeys was collected and PBMCs were purified via Ficoll density gradient centrifugation and RNA from 1 million cells was extracted according to the manufacturer’s instructions (RNeasy Mini Kit, Qiagen cat# 74104).

To generate cDNA, the SuperScript™ IV Reverse Transcriptase (cat# 18090010, Thermo Fisher Scientific) kit was used. The cDNA was cleaned using a DNA Clean and Concentrator spin column (Zymo, cat# D4033) and eluted in RNase-free water. ZAP transcript was amplified from cDNA using primer pair binding in the 5’ and 3’ ZAP-L UTR. Resulting PCR product was ligated into pCR XL TOPO Blunt vector. Sanger sequencing revealed 3 unique ZAP-L alleles. In the next step, ZAP with N-terminal HA tag was cloned into the pCG IRES BFP vector using XbaI/MluI restriction sites. All constructs were cloned using primers listed in Table 1 and their correctness was confirmed by Sanger sequencing (eurofins).

### Transduction of Jurkat cells

Vesicular stomatitis virus glycoprotein (VSV-g)-pseudotyped virus stocks were prepared by transfecting HEK293T cells with 1 μg VSV-g and 5 μg proviral construct, followed by a medium change. Supernatants were harvested 48 h later and normalized based on infectivity as measured by the TZM-bl reporter cells. One million Jurkat WT or ZAP KO cells were seeded and treated with IFN-α (200 U/ml) or left unstimulated at the time of transduction. The input virus was removed 24h later, and cells were washed in PBS. Three and six days later, supernatants were harvested and used to infect TZM-bl reporter cells in triplicates. β-Galactosidase activity was measured 2 days later using the Gal-Screen kit (Applied Biosystems) as relative light units per second using a microplate luminometer.

### Flow cytometry of infected primary cells

PHA-activated PBMCs were infected with VSV-G-pseudotyped HIV-2 or SIVsmm viruses and harvested three days post-infection for flow cytometric analysis. Cells were fixed and permeabilized (NordicMUBio, Cat#GAS-002) and then stained intracellularly for Gag (FITC conjugated; 6604665; Beckman Coulter) and ZAP (GenTex, cat# GTX120134), followed by secondary anti-rabbit AlexaFluor647 (Invitrogen, cat#A-21245). Rabbit isotype IgG antibody served as a negative control (CellSignaling cat#2729S). The MFI of infected (FITC+) cells were used to determine changes in ZAP protein levels.

### Confocal microscopy

Hela cells were seeded onto poly-Lysine coated IBIDI-slides and co-transfected with 125 ng of pCG mCherry-ZAP-L expression plasmid and 125ng of HIV-2 7312A IRES eGFP provirus or empty vector control LT1 transfection reagent. 40 h post-transfection, cells were fixed in PBS containing 2% PFA and 0.05% TritonX and stained for 30min in 1μg/ml DAPI.

### Lentiviral sensitivity to ZAP

To assess relative lentiviral sensitivity to ZAP overexpression, HEK293T ZAP KO cells (in 24-well format) were co-transfected using polyethyleneimine Max (PEI Max, Polysciences) transfection reagent with 600 ng of the indicated provirus and increasing concentrations (0, 100, 200, and 400 ng) of pCG HA-ZAP-L IRES BFP expression vector and the amount of DNA was normalized to 1 μg by adding pCG IRES GFP vector. Virus-containing supernatants were harvested 2 days later and used to infect 10.000 TZM-bl reporter cells in triplicates. β-Galactosidase activity was measured 2-3 days later using the Gal-Screen kit (Applied Biosystems) as relative light units per second using a microplate luminometer. Infectious virus yield values of each virus in the presence of ZAP were normalized to the corresponding GFP-only control.

### Lentiviral sequence analysis, CpG mapping and RNA structure prediction

Sequences of SIVsmm and HIV-2 isolates were obtained from the Los Alamos HIV sequence database (https://www.hiv.lanl.gov/content/sequence/HIV/mainpage.html). A phylogenetic tree showing the relationship between lentiviruses was generated using full-genome sequences and the ClustalW2 simple phylogeny tool (https://www.ebi.ac.uk/Tools/phylogeny/simple_phylogeny/). CpG frequency was calculated as [number of CpG]/[sequence length], whereas CpG suppression was calculated as [number of CpG * sequence length]/[number of C * number of G]. The RNAfold web server based on the latest ViennaRNA Package (Version 2.6.3; http://rna.tbi.univie.ac.at/cgi-bin/RNAWebSuite/RNAfold.cgi) was used to predict the minimum free energy (MFE) RNA structure, using the dynamic programming algorithm (41).

### Western blotting

To examine the viral protein levels under the expression of ZAP, HEK293T ZAP KO cells were co-transfected in 12-well plates with 1.25 μg of indicated provirus and 0.25 μg DNA of pCG HA-ZAP-L IRES BFP vector or empty pCG IRES BFP vector. Two days post-transfection, cells were lysed with coimmunoprecipitation (CO-IP) buffer (150 mM NaCl, 50 mM HEPES, 5 mM EDTA, 0.10% NP-40, 0.5 mM sodium orthovanadate, 0.5 mM NaF, protease inhibitor cocktail from Roche), and cell-free virions were purified by centrifugation of cell culture supernatants through a 20% sucrose cushion at 20, 800 × g for 90 min at 4°C and lysed in co-IP lysis buffer. Samples were reduced in the presence of β-mercaptoethanol by boiling at 95°C for 10 min. Proteins were separated in 4% to 12% Bis-Tris gradient acrylamide gels (Invitrogen), blotted onto a polyvinylidene difluoride (PVDF) membrane, and incubated with anti-HIV-1 Env (cat#ADP20421, CFAR), anti-HIV-2 Env (cat#ARP-1410; NIH AIDS Reagent Program), anti-HIV-1 p24 (cat#ab9071; Abcam), anti-SIVmac p27 (cat#ARP-2321; NIH AIDS Reagent Program), anti-GAPDH (Cat# 607902; BioLegend), anti-ZAP (catalogue no. GTX120134; GeneTex), anti-KHNYN (#sc-514168, Santa Cruz Biotechnology), anti-TRIM25 (cat#610570, BD), anti-Hsp90 (cat#sc7947, Santa Cruz Biotechnology) antibodies. Subsequently, blots were probed with IRDye 680RD goat anti-rabbit IgG(H+L) (catalogue no. 926-68071; LI-COR) and IRDye 800CW goat anti-mouse IgG(H+L) (catalogue no. 926-32210; LI-COR) Odyssey antibodies and scanned using a Li-Cor Odyssey reader.

### Enhanced cross-linking and immunoprecipitation (eCLIP) sequencing

HEK293T ZAP KO cells (7) (8mln) were co-transfected with 30 µg of pCG HA-ZAP-L IRES BFP and 50 µg of HIV-2 7312A or SIVsmmL1.V1 proviral DNA. After 6h, medium was changed and cells were incubated at 37°C 5%CO2 for another 40h. The culture medium was removed, and cells were washed once with ice-cold PBS followed by cross-linking (254 nm and 0.8 J/cm2 UV light) on ice. Cells were harvested into ice-cold PBS and centrifuged at 400 × g at 4°C for 5 min. The cell pellet was washed once with ice-cold PBS and frozen down at -80°C. Frozen cell pellets were lysed in 50 mM Tris-HCl pH 7.5, 150 mM NaCl, 1 mM EDTA, 1% (vol/vol) NP-40, 0.5% sodium deoxycholate, 0.25 mM TCEP [Tris(2-carboxyethyl) phosphine hydrochloride], and complete EDTA-free protease inhibitor cocktail (Roche).

Immunoprecipitation and cDNA library preparation was performed as described in the eCLIP method (PMID: 27018577), including the following modifications: immunoprecipitates were washed two times in 1 ml CLIP lysis buffer and two times in IP wash buffer (50 mM Tris-HCl pH 7.4, 300 mM NaCl, 1 mM EDTA, 1% [vol/vol] NP-40, 0.5% sodium deoxycholate, 0.25 mM TCEP), followed by two washes in 50 mM Tris-HCl pH 7.4, 1 mM EDTA, and 0.5% (vol/vol) NP-40. All other steps were carried out as described in the eCLIP method. Briefly, following cell lysis, unprotected RNA was digested with RNase I, and HA-tagged ZAP proteins were immunoprecipitated using a mouse monoclonal HA-antibody (Abcam, ab49969). Following immunoprecipitation, protein-RNA complexes were separated by SDS-PAGE and transferred onto a nitrocellulose membrane. Bands migrating at the expected size range were excised for immunoprecipitation (IP) and size-matched input (SMI) samples. Crosslinked RNA fragments were released by Proteinase K digestion and recovered RNA fragments were converted into a cDNA library for both IP and SMI samples by following the eCLIP procedure (PMID: 27018577). Sequencing was performed using the Illumina NextSeq 500 platform.

### eCLIP data analysis

Resulting paired-end sequencing libraries with read lengths of 2 × 40 nucleotides were adapter- and quality trimmed using cutadapt (v1.18). Reads shorter than 18 nt were discarded. A custom java program was applied to identify and clip the unique molecular identifier (UMI) associated with each read. The trimmed reads were then aligned to the human (hg38, Ensembl release 110) and one of the respective viral genomes (gRNA sequence based on published full genome sequences SIVsmm: GenBank KU182922.1; HIV-2: GenBank KX174311.1) using STAR (v2.7.10a) (42) with the parameters --outFilterScoreMinOverLread 0 -- outFilterMatchNminOverLread 0 --outFilterMatchNmin 0 --outFilterType Normal -- alignSoftClipAtReferenceEnds No --alignSJoverhangMin 8 --alignSJDBoverhangMin 1 -- outFilterMismatchNoverLmax 0.04 --scoreDelOpen −1 --alignIntronMin 20 --alignIntronMax 3000 --alignMatesGapMax 3000 --alignEndsType EndToEnd. PCR duplicates were removed with the UMI-aware deduplication functionality in Picard’s MarkDuplicates. Protein binding regions were predicted by using MACS2 (43) which models read coverage enrichment in an IP sample over a paired SMI control under a Poisson distribution. The analysis was performed using the parameters: -s 31 --keep-dup all --nomodel --d-min 25 --call-summits --scale-to small --shift 25 --nolambda --extsize 5 --max-gap 20 --min-length 5. The identified MACS2 peaks were further filtered by applying a one-sided Fisher’s exact test. Statistically significant enrichment was determined by calculating the odds-ratio of mapped reads within each peak against all remaining mapped reads between IP and SMI. Multiple testing correction was applied using the Benjamini-Yekutieli (44) procedure, and only peaks with an adjusted *P* value <0.05 were considered for further analysis. Additionally, the enrichment of regions of interest, such as R, U5, and U3, was calculated by using a one-sided Fisher’s exact test, following the same approach as the filtering of MACS2 peaks.

To visualize the eCLIP signal in IP relative to SMI, the relative information content of IP over SMI (45–47)was calculated as *p_i_* × log_2_(*p*_i_*/q*_i_), where *i* represents a genomic position, *p_i_* denotes the relative fraction of aligned reads covering that position in IP and *q_i_* denotes the relative fraction of aligned reads covering the same position in SMI. The relative information content was visualized using the Integrative Genome Visualization (IGV) Browser.

### Statistical analysis

Statistical analysis was performed using GraphPad Prism software. ANOVA, Mann-Whitney U-test or two-tailed Student’s t-test were used to determine statistical significance. Significant differences are indicated as: *, p < 0.05; **, p < 0.01; ***, p < 0.001. Statistical parameters are specified in the Figure legends.

## RESULTS

### SIVsmm is highly sensitive to restriction by type I IFN in primary human cells

HIV-2 originated from and is genetically closely related to SIVsmm strains but is highly divergent from HIV-1, the main causative agent of AIDS. To compare the ability of HIV-1, HIV-2 and SIVsmm to evade human intrinsic immune responses, we analysed their replication in the presence of type I IFN in primary human peripheral blood mononuclear cells (PBMCs) containing T lymphocytes (26). To ensure that the results are representative of the lentiviral groups, we selected panels of five infectious molecular clones (IMCs) of HIV-1, including four primary transmitted-founder IMCs of subtypes B and C (CH77, CH58, THRO and CH167), five HIV-2 IMCs including the patient-derived 7312A strain, and five SIVsmm strains representing diverse viral lineages (L1-L5). In the absence of IFNα, all viruses replicated efficiently, with HIV-2 strains reaching a replication peak around day 6 and SIVsmm reaching a peak at day 8-10 (Figure 1A-B). HIV-1 strains replicated very efficiently also in the presence of IFN (2-fold reduction), while HIV-2 replication was inhibited by ∼5-fold (Figure 1C). In comparison, SIVsmm strains were more strongly suppressed by type I IFN (32-fold reduction) and generally replicated poorly in its presence. Thus, SIVsmm exhibits lower replication fitness and immune evasion than HIV-1 and HIV-2, suggesting suboptimal adaptation to type I IFN-induced antiviral restriction factors.

**Figure 1.**
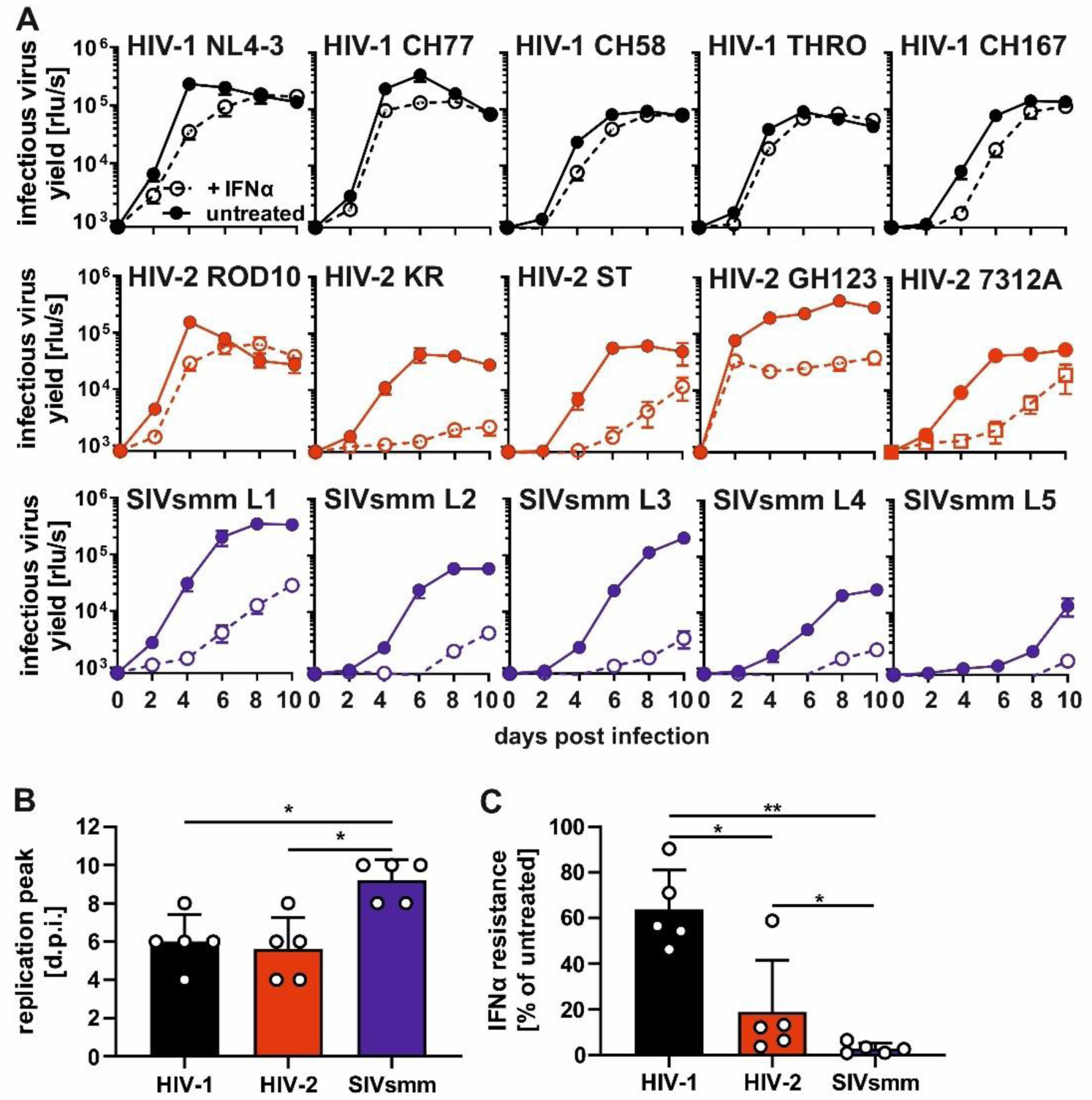
Replication fitness and type I IFN sensitivity of HIV-1, HIV-2 and SIVsmm in primary human cells. (A) Replication kinetics of indicated lentiviral strain in pre-activated and IFNα-treated (500u/ml; dotted line) or untreated (solid line) human PBMCs up to 10 days post-infection. Infectious virus yield was quantified using a TZM-bl reporter assay. Shown is the mean of 3 individual donors, each tested in triplicates of infection, +/- SEM. (B) Average time needed to reach a replication peak of each virus group based on (A). Mean + SD. (C) Resistance to IFNα treatment based on (A). Mean + SD. *, P < 0.05; **, P < 0.01; calculated by Mann–Whitney test.

### HIV-2 evolved ZAP resistance

ZAP is a broadly acting antiviral restriction factor that forms an antiviral complex with TRIM25 ubiquitin ligase and endoribonuclease KHNYN to degrade CpG-containing viral transcripts. Their expression is upregulated by IFN (Figure S1A) (3, 4), which suggests that ZAP antiviral complex might contribute to the restriction of SIVsmm in human PBMCs.

To determine the contribution of ZAP to IFN susceptibility of HIV-2 and SIVsmm, we used Jurkat CCR5 CRISPR Cas9 ZAP KO cells (22). This cell line was selected due to its resemblance to human T lymphocytes, the natural target cells of HIV. The infection of control and ZAP KO Jurkat cells with HIV-1, HIV-2 and SIVsmm resulted in a detectable virus production of all strains (Figure 2A, S1B). ZAP KO modestly increased infectious HIV-1 production by 50-100% in the absence and presence of IFNα, while HIV-2 was unaffected by ZAP (Figure 2B). In contrast, ZAP KO enhanced SIVsmm infection by 3-fold in the absence of IFNα and up to 5-fold in the presence of IFNα treatment. Thus, ZAP restricts SIVsmm more efficiently than HIV-1 and HIV-2 in human T cells.

**Figure 2.**
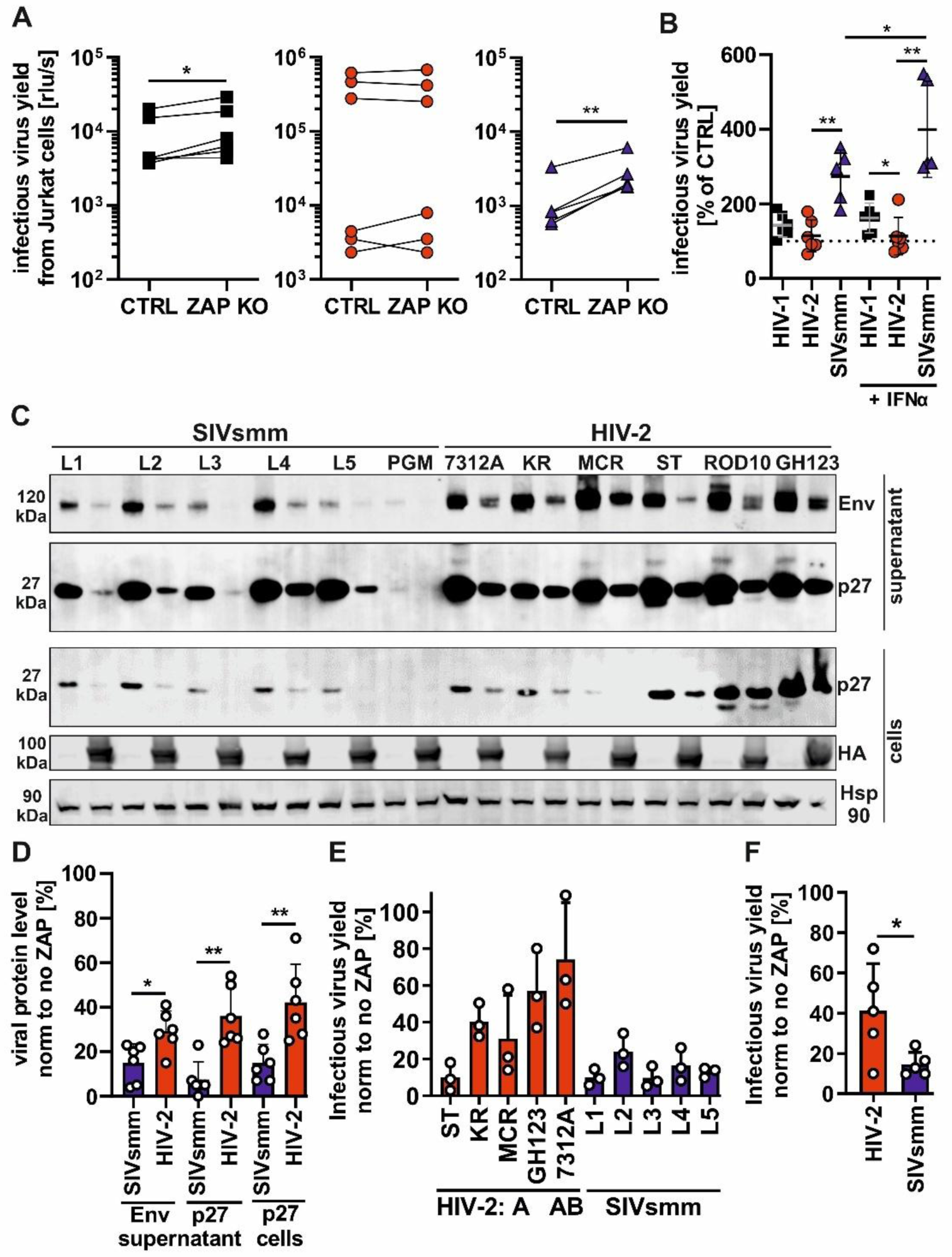
Inhibition of HIV-2 and SIVsmm by ZAP. (A) HIV-1 (black square), HIV-2 (red circle) and SIVsmm (blue triangle) infectious virus production in parental (CTRL) or ZAP knock out (KO) CCR5-expressing Jurkat cells 6 days after transduction. (B) Infectious virus yield % calculated by dividing virus yield in the absence (KO) and presence (CTRL) of ZAP, in the absence and presence of 500u/ml IFNα treatment. (C) western blot of HEK293T ZAP KO cells and their supernatants following the transfection of empty BFP vector control or HA-tagged ZAP and indicated infectious molecular clone of SIVsmm or HIV-2, and (D) its quantification. Hsp90 serves as loading control. (E) Infectious virus yield of HIV-2 and SIVsmm produced from HEK293T ZAP KO cells co-transfected with indicated infectious molecular clone and human HA-tagged ZAP. Values were normalized to infectious virus produced in the absence of ZAP expression. (F) Average infectious virus produced from HEK293T ZAP KO cells in the presence of ZAP overexpression based on (E). N=3 + SD. *, P < 0.05; **, P < 0.01; calculated using Student’s t-test.

To ensure that these results were not biased by relatively low replication of SIVsmm strains in this system, we performed additional overexpression experiments in HEK293T ZAP KO cells (7). For this, we used an expression plasmid encoding the long isoform of ZAP (ZAP-L), which is the most active isoform of ZAP (2, 48, 49). Transfection of SIVsmm IMCs resulted in similar or slightly lower viral protein expression and infectious virus yield than HIV-2 (Figure S1C), in agreement with previous studies data (13, 26). On average, ZAP reduced the p27 and Env protein production of SIVsmm by ∼90% but only by ∼65% in the case of HIV-2 (Figures 2C-D). HIV-2 strains differed in ZAP sensitivity from being highly resistant (HIV-2 7312A, directly isolated from patient PBMCs) to being as sensitive as SIVsmm (HIV-2 ST, passaged in cell lines) (Figure 2E). In general, SIVsmm virus production was on average 3.5 times more inhibited by ZAP than that of HIV-2 (Figure 2F). Altogether, these results clearly suggest that HIV-2 evolved increased ZAP resistance during its adaptation to humans.

### HIV-2 evades ZAP despite higher CpG content in a species-independent manner

CpG suppression plays a key role in the evasion of ZAP restriction of many viruses (1, 2, 7, 50). For the set of HIV-2 and SIVsmm IMCs analysed, however, high CpG numbers did not correlate with increased ZAP sensitivity and even showed the opposite trend (Figure 3A). On average, HIV-2 strains contained about two times more CpGs than HIV-1 (Figure 3B). Extended analysis of the 72 available full viral genome sequences from the Los Alamos HIV database confirmed that epidemic HIV-2 A and B strains show significantly higher CpGs frequencies than SIVsmm (Figure 3C). In contrast, the CpG frequencies in rare HIV-2 F-I did not differ significantly from those of SIVsmm. Thus, epidemic HIV-2 strains evolved ZAP resistance despite an overall increase in CpG content. We have previously shown that the sensitivity of HIV-1 to ZAP is determined by the number of CpGs in the first 600 bp of the *env* ORF rather than the whole genome (13). Thus, we compared the distribution of CpGs across HIV-2 and SIVsmm genomes to identify site-specific differences in CpG content (Figure 3D). HIV-2 contained an increased number of CpGs throughout the genome. The location of CpGs in HIV-2 7312A substantially overlapped with those found in SIVsmmL1 (Figure 3E). Furthermore, HIV-2 did not decrease ZAP expression levels or affect subcellular localization (Figure S2A-C) suggesting that it evades rather than directly counteracts ZAP.

**Figure 3.**
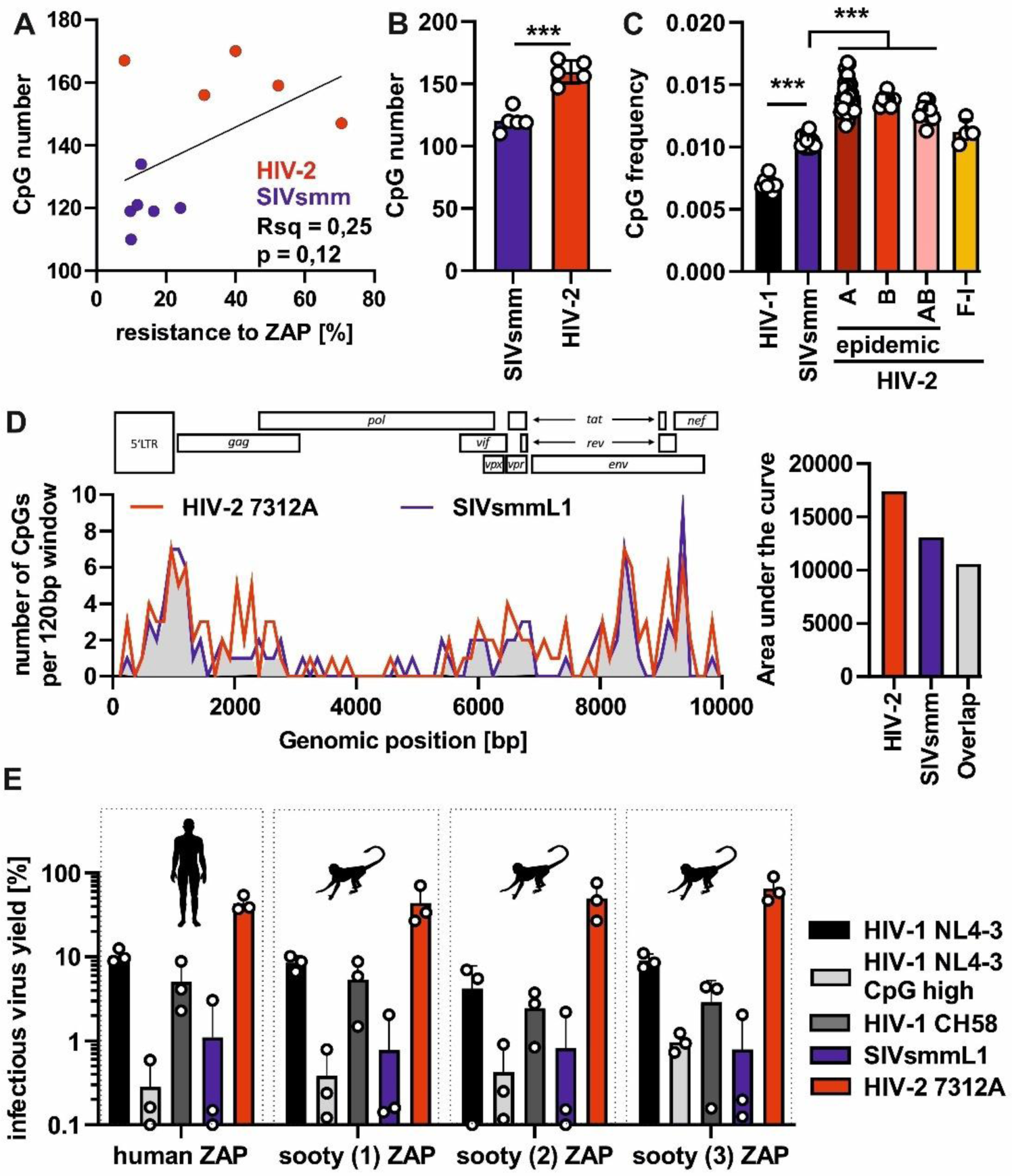
CpG number and distribution in HIV-2 and SIVsmm genomes and the antiviral activity of human and sooty ZAP orthologues. (A) Correlation between CpG number in HIV-2 (red) and SIVsmm (blue) strain genomes and their resistance to ZAP overexpression (related to Figure 2E). (B) Average CpG number in the tested HIV-1, HIV-2 and SIVsmm strains and (C) average CpG frequency in HIV-1, SIVsmm and HIV-2 (groups A-I) full-genome sequences obtained from the Los Alamos HIV sequence database. (D) Left: distribution of CpGs in the aligned HIV-2 7312A (red) and SIVsmm L1 (blue) genomes and overlap in CpG positions (grey). ORF position in the HIV-2 genome is shown for reference above the graph. Right: Quantification of the area under the curve for HIV-2, SIVsmm and overlapping peaks. (E) Infectious virus yield of HIV-1, HIV-2 and SIVsmm co-transfected with 0.4 µg DNA of human and sooty mangabey ZAP in ZAP KO HEK293T cells. N=3 + SD. ***, P < 0.001, calculated using the Mann–Whitney test.

Single-stranded RNA binding domains (RBD) of ZAP orthologues from different vertebrate species exhibit varying degrees of activity and CpG-specificity (10). To determine if differences between human and sooty mangabeys ZAP activity could explain the high sensitivity of SIVsmm, we amplified ZAP from the cDNA of captive sooty mangabeys. We obtained three ZAP variants and cloned them into an expression vector. The otherwise highly conserved RNA binding domain (RBD) of sooty mangabey ZAP differed at three positions from human ZAP (Figure S3A). To determine the potential impact of these mutations on antiviral activity, we co-transfected HEK293T ZAP KO cells with ZAP and HIV-1, HIV-2 or SIVsmm proviruses. Sooty mangabey ZAP strongly inhibited the control CpG-enriched HIV-1 mutant virus and restricted HIV-1 and SIVsmm as efficiently as human ZAP, while HIV-2 7312A was generally less affected by either orthologue (Figure 3E). Thus, sooty mangabey ZAP is as active as its human orthologue and HIV-2 evolved resistance to both, which might have reduced the selective pressure driving CpG suppression in its RNA genome.

### HIV-2 RNA is bound by ZAP but resistant to its antiviral effects due to changes in the 3’ half of the genome

RNA binding is essential for the antiviral activity of ZAP (3, 48). To compare ZAP binding events across the HIV-2 and SIVsmm RNA, we performed enhanced cross-linking and immunoprecipitation in combination with RNA sequencing analysis (eCLIP-seq) in HEK293T ZAP KO cells co-transfected with HIV-2 or SIVsmm proviruses and ZAP. We normalized the signal observed in ZAP immunoprecipitations relative to a size-matched input control and observed seven significantly enriched peak regions (FDR <0.05, log 2 fold change > 0.5; Table 2), all of which contained at least one CpG (Figure 4A). In the case of HIV-2 RNA, we identified nine significantly enriched ZAP-binding sites (FDR <0.05, log2 fold change > 0.5; Table 2) of which five overlapped with CpG positions (Figure 4B). In both viruses, most ZAP binding sites localized to the 3’ genome half (downstream of the *vpr* gene). To analyse the impact of this region on ZAP resistance, we generated chimeras between the resistant HIV-2 7312A and sensitive SIVsmmL1. To avoid changes within the viral ORFs, we selected the *vpx/vpr* gene border (Figure 4C) to generate IMCs carrying 5’ and 3’ genome halves of HIV-2 or SIVsmm, respectively. Chimeric virus containing the 3’ half of the HIV-2 genome was more resistant to ZAP overexpression than the virus containing the 3’ half of SIVsmm (Figures 4D, S4A). Thus, ZAP sensitivity is determined by the 3’ viral genome half which contains most of the ZAP binding sites.

**Figure 4.**
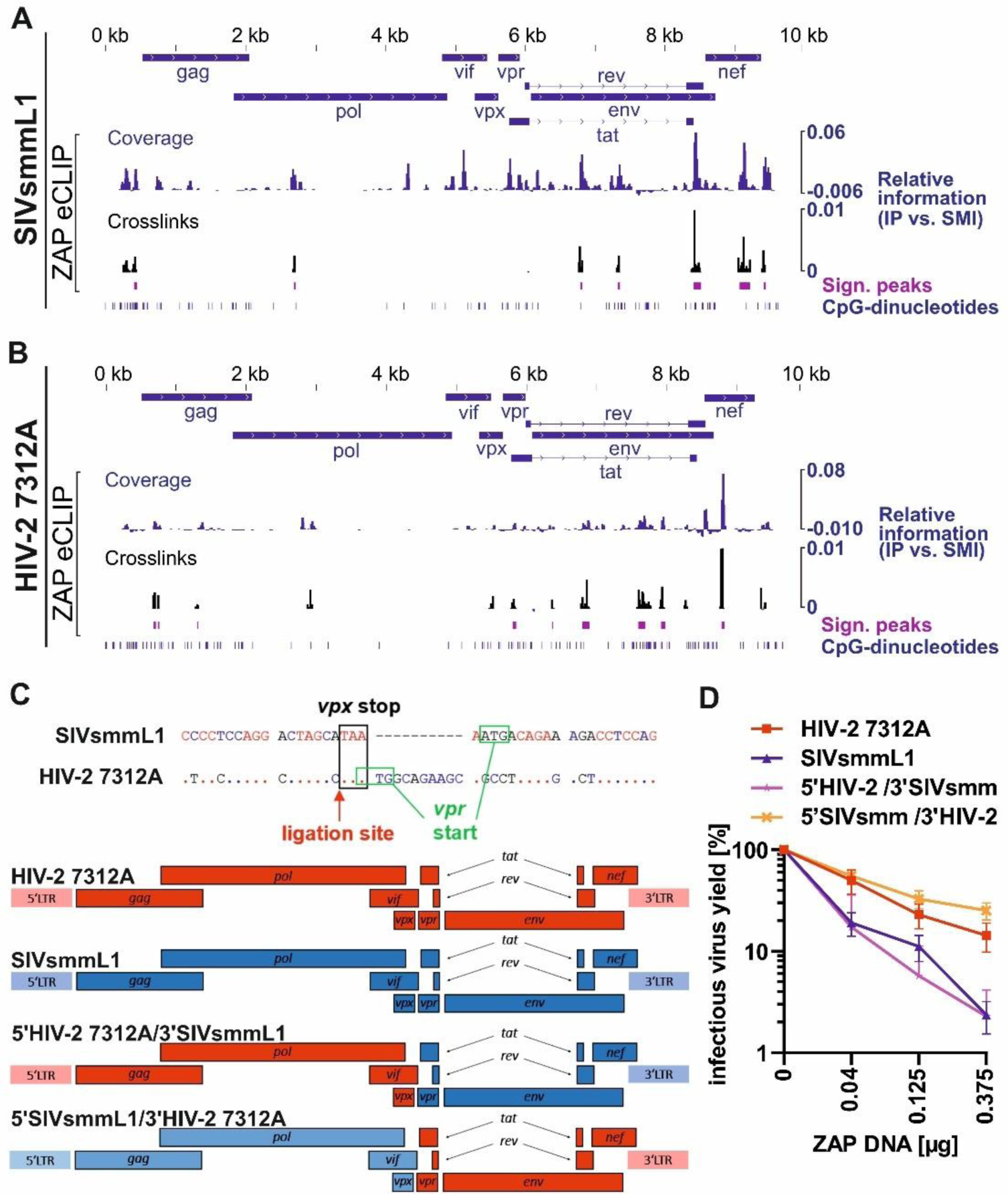
Analysis of ZAP binding regions in HIV-2 and SIVsmm RNA and mapping of the determinants of ZAP resistance. (A) Alignment of HA-ZAP eCLIP data to SIVsmmL1 and (B) HIV-2 7312A genomic RNA. Relative coverage information in IP vs. size-matched input (SMI) is calculated at each position and displayed along the viral genome (blue). ZAP binding sites significantly enriched relative to SMI are shown in dark pink. For each significantly enriched ZAP binding site, crosslinking events are extracted and shown in black (relative information in IP vs. SMI). CpG-dinucleotides are displayed in dark blue. (C) Position of ligation of the half HIV-2/ half SIVsmm virus chimeras and schematic of generated chimeric proviral constructs showing HIV-2 (red) and SIVsmm (blue) fragments. (D) Sensitivity of those chimeras to ZAP overexpression in HEK293T ZAP KO cells. N=3+ SD.

**Table 2:**
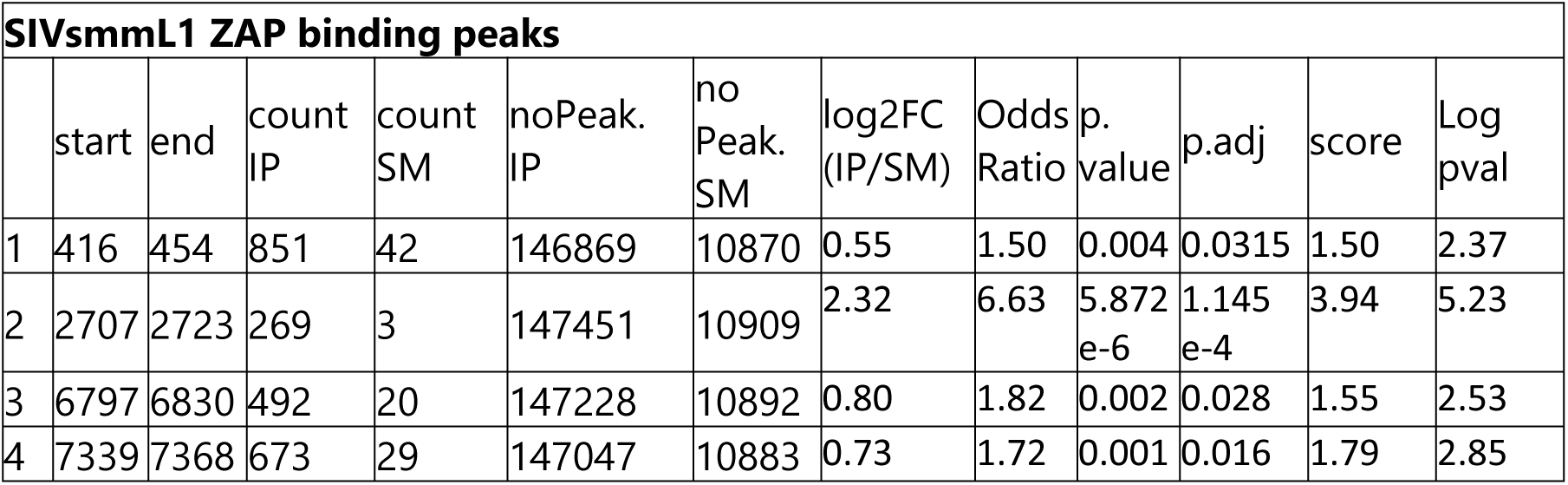
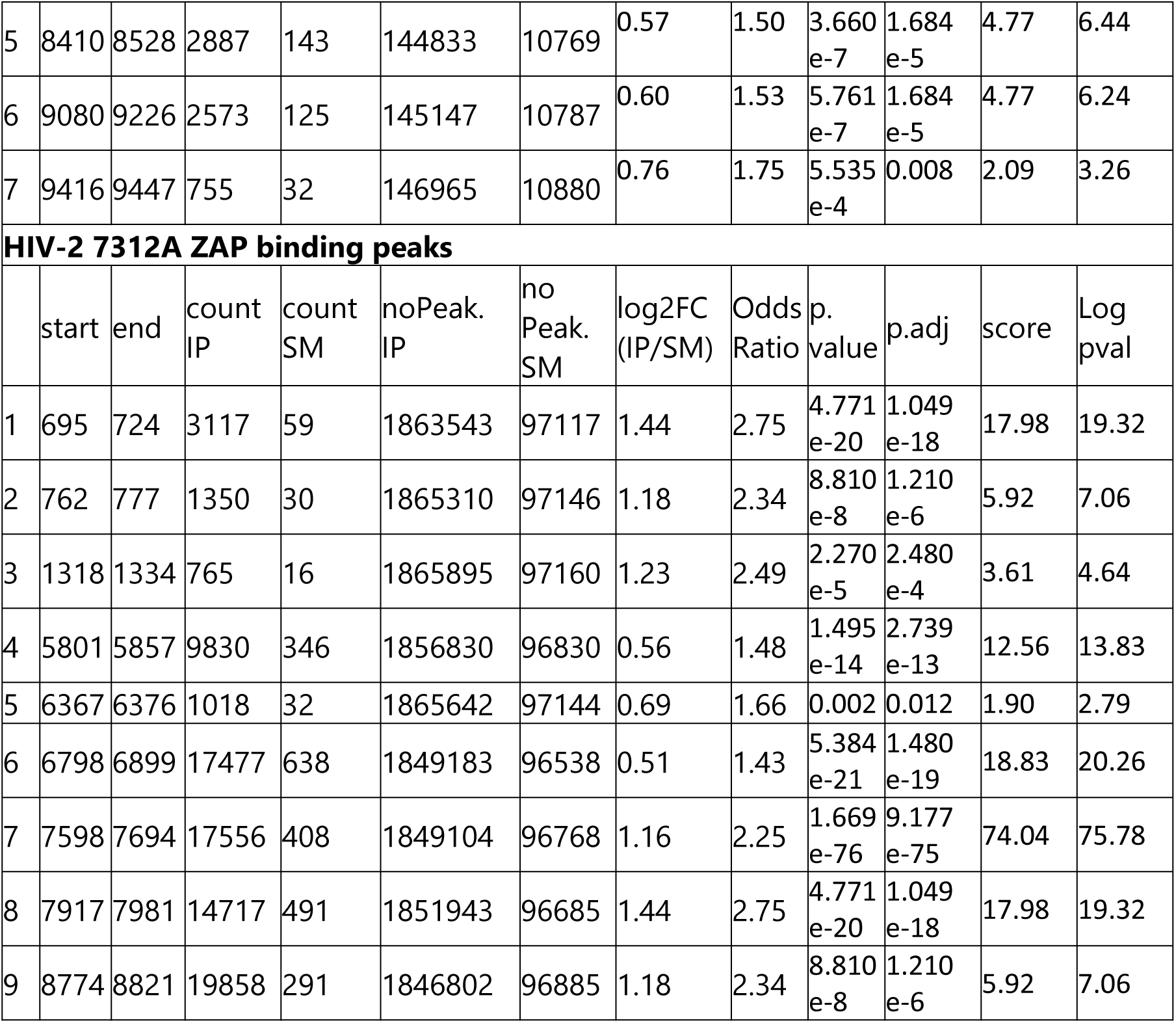
Significant peaks identified by eCLIP.

| SIVsmmL1 ZAP binding peaks |  |  |  |  |  |  |  |  |  |  |  |  |
| --- | --- | --- | --- | --- | --- | --- | --- | --- | --- | --- | --- | --- |
|  | start | end | count<br>IP | count<br>SM | noPeak.<br>IP | no<br>Peak.<br>SM | log2FC<br>(IP/SM) | Odds<br>Ratio | p.<br>value | p.adj | score | Log<br>pval |
| 1 | 416 | 454 | 851 | 42 | 146869 | 10870 | 0.55 | 1.50 | 0.004 | 0.0315 | 1.50 | 2.37 |
| 2 | 2707 | 2723 | 269 | 3 | 147451 | 10909 | 2.32 | 6.63 | 5.872<br>e-6 | 1.145<br>e-4 | 3.94 | 5.23 |
| 3 | 6797 | 6830 | 492 | 20 | 147228 | 10892 | 0.80 | 1.82 | 0.002 | 0.028 | 1.55 | 2.53 |
| 4 | 7339 | 7368 | 673 | 29 | 147047 | 10883 | 0.73 | 1.72 | 0.001 | 0.016 | 1.79 | 2.85 |
| 5 | 8410 | 8528 | 2887 | 143 | 144833 | 10769 | 0.57 | 1.50 | 3.660<br>e-7 | 1.684<br>e-5 | 4.77 | 6.44 |
| 6 | 9080 | 9226 | 2573 | 125 | 145147 | 10787 | 0.60 | 1.53 | 5.761<br>e-7 | 1.684<br>e-5 | 4.77 | 6.24 |
| 7 | 9416 | 9447 | 755 | 32 | 146965 | 10880 | 0.76 | 1.75 | 5.535<br>e-4 | 0.008 | 2.09 | 3.26 |
| **HIV-2 7312A ZAP binding peaks** |  |  |  |  |  |  |  |  |  |  |  |  |
|  | start | end | count<br>IP | count<br>SM | noPeak.<br>IP | no<br>Peak.<br>SM | log2FC<br>(IP/SM) | Odds<br>Ratio | p.<br>value | p.adj | score | Log<br>pval |
| 1 | 695 | 724 | 3117 | 59 | 1863543 | 97117 | 1.44 | 2.75 | 4.771<br>e-20 | 1.049<br>e-18 | 17.98 | 19.32 |
| 2 | 762 | 777 | 1350 | 30 | 1865310 | 97146 | 1.18 | 2.34 | 8.810<br>e-8 | 1.210<br>e-6 | 5.92 | 7.06 |
| 3 | 1318 | 1334 | 765 | 16 | 1865895 | 97160 | 1.23 | 2.49 | 2.270<br>e-5 | 2.480<br>e-4 | 3.61 | 4.64 |
| 4 | 5801 | 5857 | 9830 | 346 | 1856830 | 96830 | 0.56 | 1.48 | 1.495<br>e-14 | 2.739<br>e-13 | 12.56 | 13.83 |
| 5 | 6367 | 6376 | 1018 | 32 | 1865642 | 97144 | 0.69 | 1.66 | 0.002 | 0.012 | 1.90 | 2.79 |
| 6 | 6798 | 6899 | 17477 | 638 | 1849183 | 96538 | 0.51 | 1.43 | 5.384<br>e-21 | 1.480<br>e-19 | 18.83 | 20.26 |
| 7 | 7598 | 7694 | 17556 | 408 | 1849104 | 96768 | 1.16 | 2.25 | 1.669<br>e-76 | 9.177<br>e-75 | 74.04 | 75.78 |
| 8 | 7917 | 7981 | 14717 | 491 | 1851943 | 96685 | 1.44 | 2.75 | 4.771<br>e-20 | 1.049<br>e-18 | 17.98 | 19.32 |
| 9 | 8774 | 8821 | 19858 | 291 | 1846802 | 96885 | 1.18 | 2.34 | 8.810<br>e-8 | 1.210<br>e-6 | 5.92 | 7.06 |

### HIV-2 evades ZAP restriction by an *env*-independent mechanism

The largest ORF found in the 3’ half of the HIV-2 and SIVsmm genomes encodes for the Env protein. We have previously shown that CpG content of the *env* encoding region determines ZAP sensitivity of HIV-1 (13). However, CpG levels in the *env* genes of HIV-2 and SIVsmm did not correlate with ZAP restriction (Figure S4B). To determine if the *env* coding region contributed to the difference in ZAP sensitivity of HIV-2 and SIVsmm, we utilized the published HIV-2 7312A recombinants encoding different SIVsmm Env ectodomains (16) (Figure 5A). HIV-2 WT and all chimeras were resistant to ZAP overexpression, while SIVsmmL1 was strongly inhibited (Figure 5B). To additionally determine the contribution of *env* regions encoding for the transmembrane protein domain, we utilized the bicistonic *env* IRES eGFP reporter constructs (Figure 5C). In this system, ZAP targeting of the *env* region in the RNA transcript reduces GFP expression. Comparison of GFP signal of the control HIV-2 7312A-IRES-GFP and SIVsmm PGM-IRES-GFP reporter viruses in the absence and presence ZAP, showed significant inhibition of SIVsmm but not HIV-2 7312A, indicating that the IRES eGFP cassette itself did not change ZAP phenotype (Figure 5D, left). In contrast, ZAP decreased GFP signal of both HIV-2 and SIVsmm *env*-IRES-eGFP reporters, and there was no significant difference between them (Figure 5D, right). These results show that the determinants of ZAP sensitivity of the HIV-2/SIVsmm lineage differ from that of HIV-1 and are *env*-independent.

**Figure 5.**
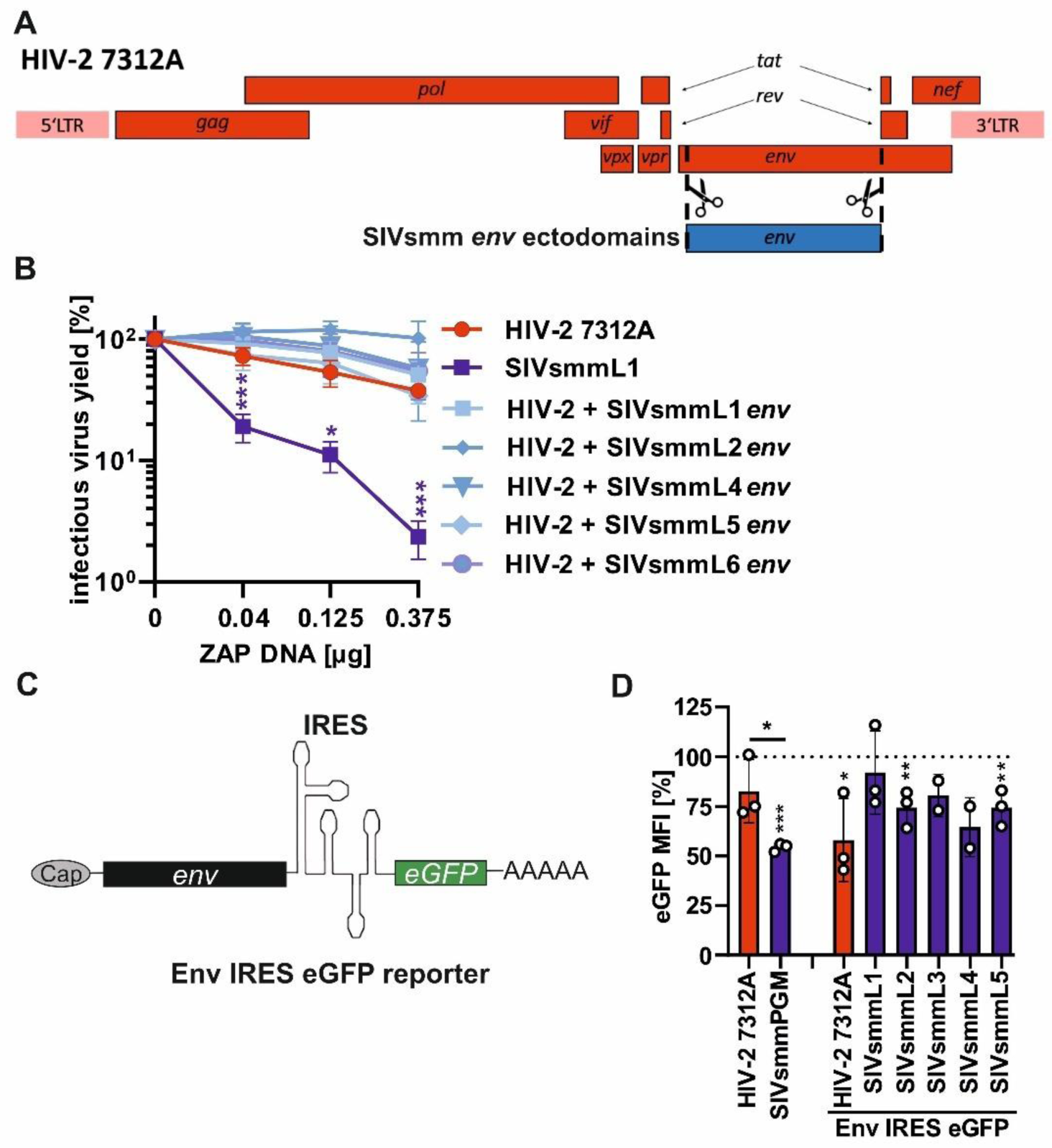
Impact of ZAP on HIV-2 Env chimeras with SIVsmm Env ectodomains and viral env transcripts. (A) Schematic of HIV-2 7312A based Env ectodomain chimeras and (B) their sensitivity to ZAP overexpression compared to wild type HIV-2 7312A and SIVsmmL1 strains. (C) Schematic of env IRES eGFP reporter and (D) flow cytometry results showing eGFP mean fluorescence intensity in HEK293T ZAP KO cells co-transfected with ZAP or ZAP mutant without RNA binding domain (ZAP delta RBD; negative control; 100%) and either IRES eGFP reporter viruses or pCG Env IRES eGFP reporter variants. (D) N=3 + SD. *, p < 0.05; **, p < 0.01; ***, p < 0.001, calculated using Student’s t-test.

### HIV-2’s *nef* encoding region contributes to ZAP resistance

The accessory proteins of HIV are known to antagonize several antiviral restriction factors (51). Therefore, we tested whether the HIV-2 accessory Vpr and Nef proteins, encoded by the 3’ genome half, counteract ZAP. HIV-2 7312A with premature stop codons in *vpr* or *nef* remained resistant to ZAP (Figure 6A), indicating that these proteins do not act as ZAP antagonists.

**Figure 6.**
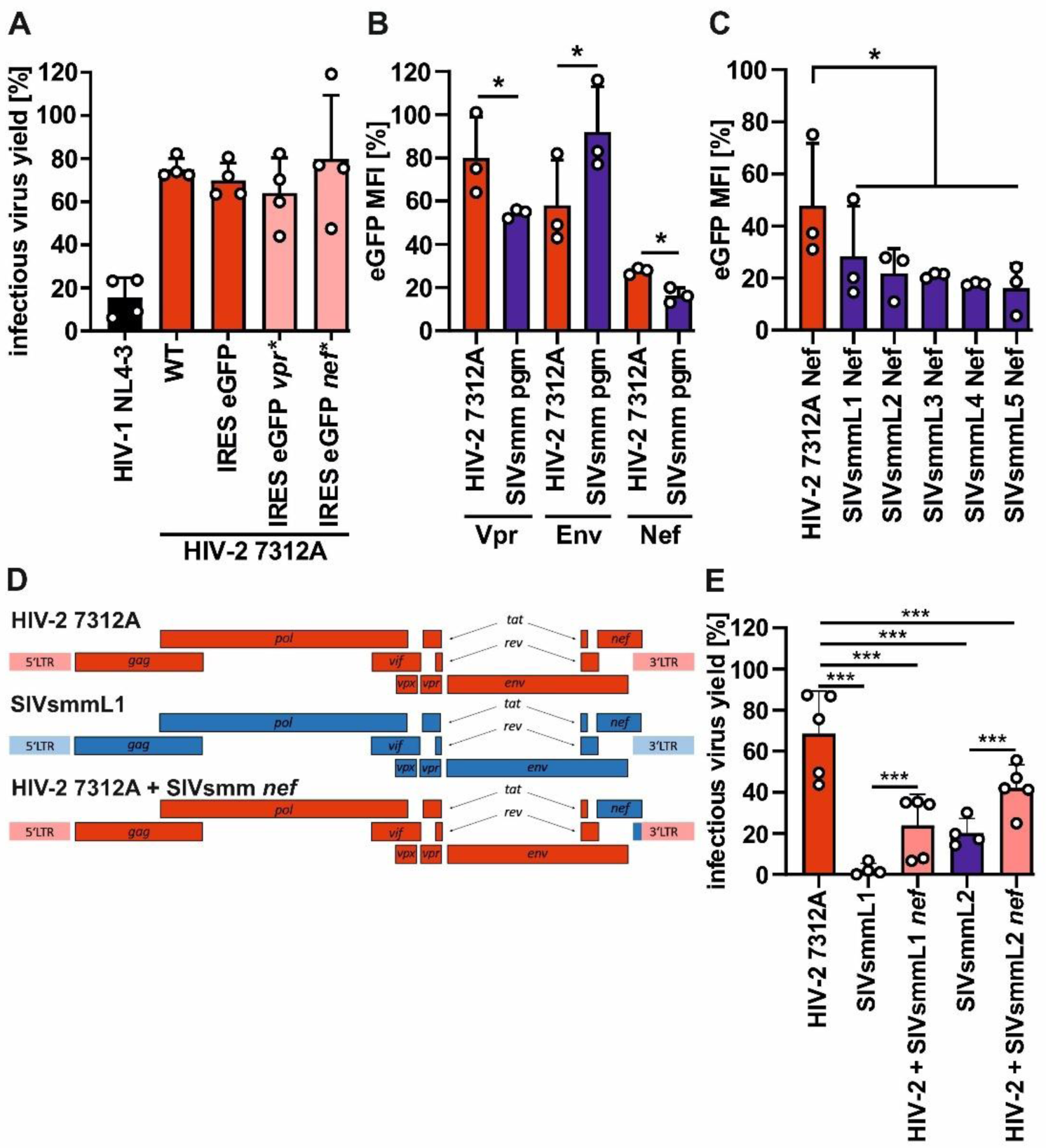
Mapping of determinants of ZAP resistance using chimeric HIV-2/SIVsmm clones. (A) Effect of ZAP overexpression (0.3 µg) on HIV-2 7312A wild type, reporter and mutants containing premature stop codons (*) in vpr and nef ORFs in HEK293T ZAP KO cells. (B) eGFP mean fluorescence intensity in HEK293T ZAP KO cells co-transfected with ZAP or ZAP delta RBD (negative control; 100%), and pCG IRES eGFP vectors containing the vpr, env or nef ORF from HIV-2 7312A or SIVsmm pgm or (C) nefs from HIV-2 7312A or SIVsmm L1-L5 (F). (D) Schematic of nef chimeric proviral constructs showing HIV-2 (red) and SIVsmm (blue) fragments and (E) Sensitivity of chimeric viruses (light pink) to ZAP overexpression (0.4 µg) relative to WT HIV-2 (red) and SIVsmm (blue) strains. N=3-5 + SD. *, P < 0.05; **, P < 0.01; ***, P < 0.001, calculated using Student’s t-test.

To examine whether HIV-2 and SIVsmm *vpr* and/or *nef* encoding regions contribute to ZAP sensitivity, we utilized an extended panel of bicistronic IRES eGFP reporters. ZAP reduced eGFP expression from mRNAs containing SIVsmm *vpr* and *nef* genes significantly more efficiently than the corresponding HIV-2 constructs (Figure 6B). The reporter containing SIVsmm *nef* ORF was more efficiently inhibited (85% decrease) by ZAP than that containing *vpr* (45% decrease). Further analyses demonstrated that bicistronic IRES-eGFP constructs containing *nef* alleles from diverse SIVsmm variants are also more sensitive to ZAP restriction than the construct containing HIV-2 7312A *nef* (Figure 6C).

To determine the impact of the *nef* coding region on viral resistance to ZAP, we generated additional chimeric HIV-2 7312A/SIVsmm viruses (Figure 6D). Unfortunately, the SIVsmmL1 chimera containing HIV-2 *nef* genes were poorly infectious precluding meaningful analyses (Fig. S4A), therefore two HIV-2-based constructs were analysed instead, one carrying the *nef* of SIVsmmL1 and the second with the *nef* of SIVsmmL2. Introduction of SIVsmm *nef* region increased HIV-2’s sensitivity to ZAP by 30-50% (Figure 6E). These results show that the *nef* ORF is an important determinant of the differential ZAP susceptibility of HIV-2 and SIVsmm.

### The *nef*/U3 LTR overlap region determines ZAP resistance of HIV-2

In the SIVsmm/HIV-2 lineage, the *nef* ORF overlaps with the U3 region of 3’LTR by ∼350bp (Figure S5). The U3 region is duplicated at 5’ and 3’ ends in the proviral (DNA) form of the viral genome, but is only found at the 3’ end of the genomic and subgenomic RNA (52) (53). To define a potential contribution of this region to ZAP sensitivity, we swapped the entire viral LTRs or its individual U5, R and U3 parts between HIV-2 and SIVsmm L1 (Figure 7A). The exchange of the LTRs completely reversed the ZAP phenotypes of HIV-2 and SIVsmm (Figure 6B). However, it also reduced basal infectious virus production by one to two orders of magnitude even in the absence of ZAP (Figure S6A). Exchange of only the 3’ U3 or U5 regions also significantly attenuated HIV-2 7312 by 1.5 and 3.5-fold correspondingly, while mutation of the 3’R region had no significant impact (Figure S6B). The analysis of these 3’LTR chimeras showed that the exchange of the U3 region sensitizes HIV-2 to ZAP (Figure 7C). Thus, the U3 region which partially overlaps with the *nef* ORF is a major determinant of HIV-2’s ZAP resistance.

**Figure 7.**
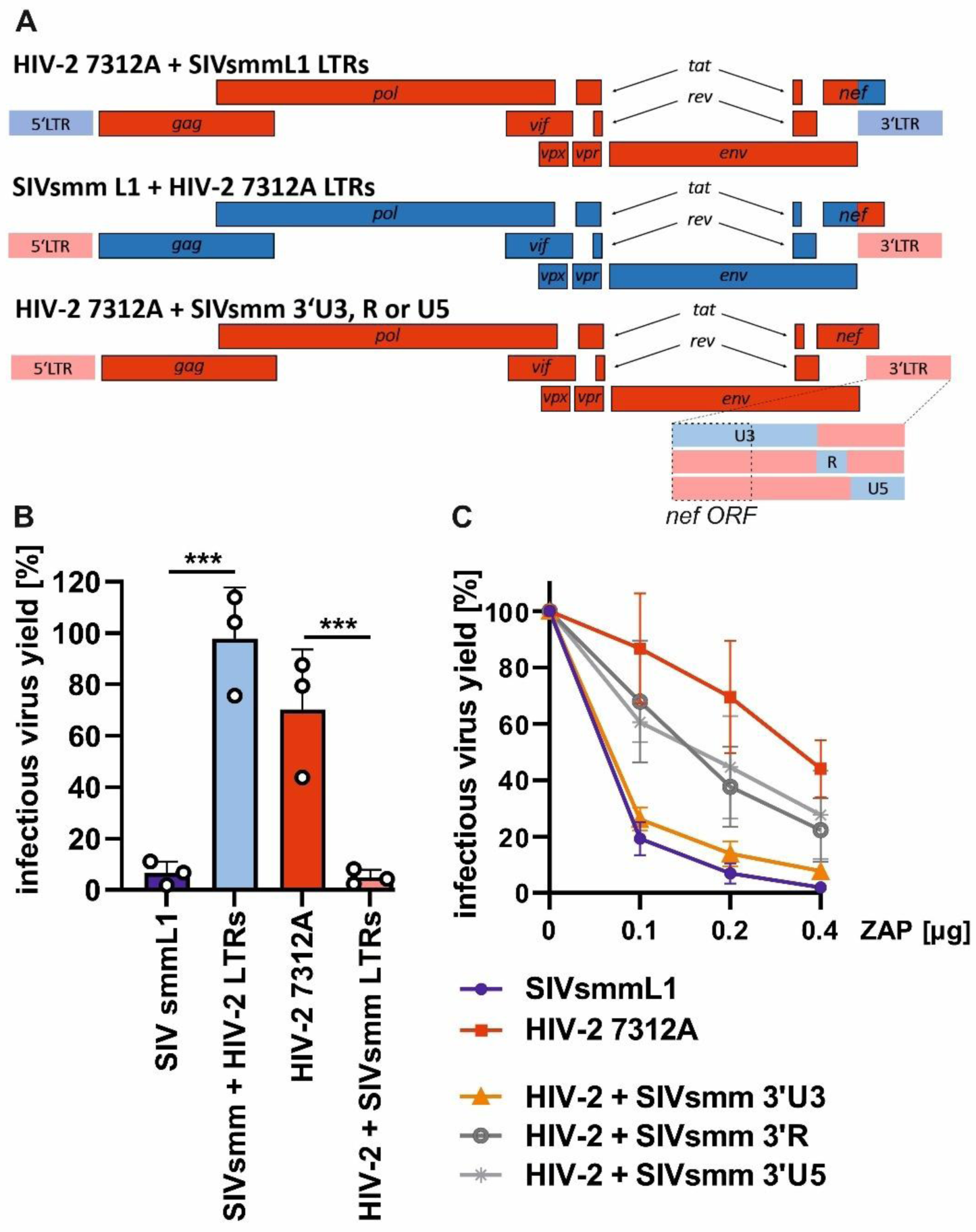
Impact of LTRs on HIV-2’s resistance to ZAP. (A) Schematic of the generated chimeric proviruses in which either both the 5’ and 3’ LTRs are exchanged, or only the U3, R or U5 region of the 3’LTR. SIVsmm L1 regions are shown in blue while HIV-2 7312A regions are shown in red. (B) Sensitivity of 5’+3’ LTR chimeras and (C) 3’ LTR U3, R and U5 chimeras to ZAP overexpression in HEK293T ZAP KO cells. N=3 + SD. *, P < 0.05; ***, P < 0.001; calculated using Student’s t-test.

### An adaptation in the U3 region of HIV-2 that mediates ZAP resistance promotes viral fitness in primary human T cells

The iCLIP analysis of viral RNA regions bound by ZAP (Figure 4A-B) identified 2 unique binding regions within the SIVsmm L1 U3 (Figure 8A). In comparison, this region was not bound in HIV-2 7312A RNA (Figure 8B). Sequence alignments revealed three CpG sites located in SIVsmmL1 but not in the HIV-2 7312A U3, two of which were found in the proximity of ZAP binding site (Figure 8C). To test whether any of these sites was specifically targeted by ZAP and responsible for differential restriction of both viruses, we generated single mutants that were depleted (SIVsmm) or enriched (HIV-2) in CpGs at the identified sites. The introduction or removal of these CpGs did not have a significant effect on viral infectivity in the absence or presence of ZAP (Figure 8D, S6C).

**Figure 8.**
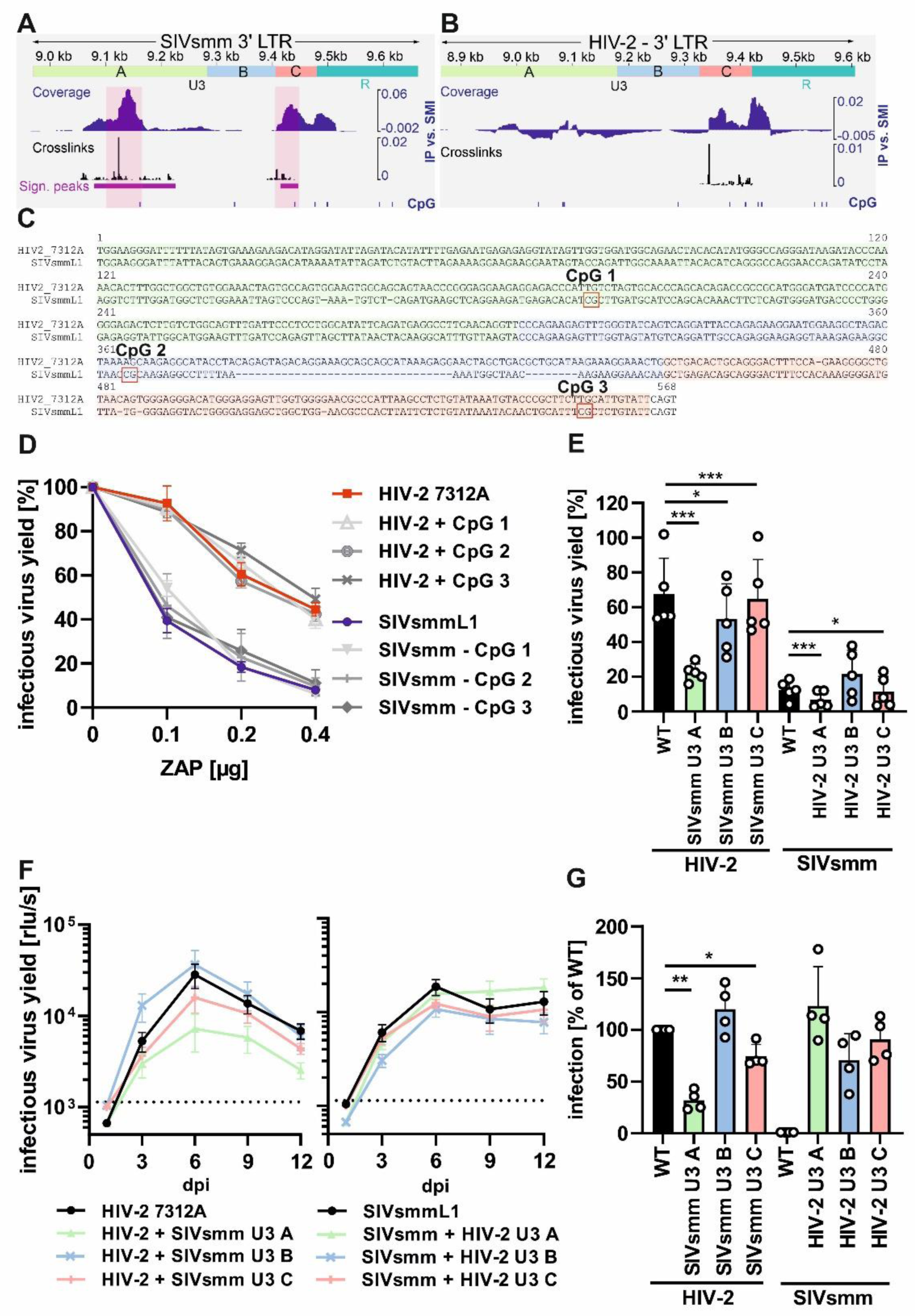
Mapping of U3 region responsible for HIV-2’s resistance to ZAP and its impact on viral fitness. (A) ZAP eCLIP data aligned to SIVsmmL1 and (B) HIV-2 7312A 3’ LTR region (based on Figure 4A-B). (C) Alignment of the HIV-2 7312A and SIVsmmL1 U3 sequences showing the location of CpG dinucleotides that are exclusively found in SIVsmmL1 (CpG 1, 2 and 3) and U3 regions A (green), B (blue) and C (pink). (D) Infectious virus yield of HIV-2 and SIVsmm CpG mutants or (E) chimeric viruses carrying exchanges in the 3’ U3 region A, B or C, in HEK293T ZAP KO cells transfected with ZAP. N=3-5 +SD. (F) Replication kinetics of chimeric HIV-2 and SIVsmm viruses carrying exchanges in the 3’ U3 region A, B or C in human PBMCs over 12 days. (G) Normalized infectious virus yield of indicated U3 mutants as compared to WT HIV-2 7312A or SIVsmmL1 (100%) based on area under the curve analysis of data shown in panel D. Mean of 4 independent donors + SD.

To map the changes that contribute to resistance to ZAP, we divided the U3 into 3 regions (Figure 8C; region A – green, B – blue and C – pink). Regions A and C contained the ZAP binding sites (ZBR) identified by iCLIP in SIVsmm U3. Region B contained no ZBR and was the most divergent region between HIV-2 and SIVsmms (Figure S7A). Previous SHAPE structure probing of the SIVmac239 RNA (54), a virus closely related and derived from SIVsmm, indicated that region B can fold a stable hairpin structure (Figure S7B), which according to our *in silico* RNA fold predictions does not form in HIV-2 (Figure S7C-D-E). To determine if changes within these regions contribute to ZAP resistance, we generated HIV-2/SIVsmm chimeras with exchanges in the U3 region A, B and C (Figure S8A-B). The introduction of SIVsmm U3 region A into HIV-2 had the most profound effect on ZAP phenotype, leading to a 3-fold decrease in ZAP resistance, while the reverse chimera in SIVsmm backbone remained highly ZAP sensitive (Figure 8C). HIV-2 with U3 region B of SIVsmm also showed a minor loss of ZAP resistance, while the corresponding SIVsmm region B mutant a minor increase in resistance (Figure 8E). In comparison, the exchange of the U3 region C had no effect on either virus. Thus, U3 region A is the main determinant of HIV-2’s ZAP resistance.

To determine if the identified U3 region A associated with ZAP resistance contributes to HIV-2 fitness in primary human cells, we examined the replication of the chimeric HIV-2 and SIVsmm viruses in pre-activated human PBMCs (Figure 8F). The exchange of U3 region A reduced the replication of HIV-2 7312A by 3.5-fold, while the exchange of the other two U3 regions had no (region B) or only modest (1.3-fold; region C) effect (Figure 8G). The corresponding SIVsmmL1-based chimeras showed no significant differences in replication.

Together, these results show that ZAP binds CpG sites in SIVsmm U3 LTR and that HIV-2 7312A evolved changes in the U3 region promoting ZAP escape and replication fitness through a mechanism independent of CpG-depletion.

## DISCUSSION

ZAP restricts the replication of many RNA and DNA viruses, often in an IFN-dependent manner (1). The ability of ZAP to preferentially target foreign RNA transcripts relies on its selective binding to CpG dinucleotides (7) which represent the least abundant type of dinucleotide in the vertebrate genomes (55). HIV-1 and many other RNA viruses strongly suppresses their CpG content and thus at least partially escape ZAP’s activity (7, 50, 56–59). Here, we found that epidemic HIV-2 evolved resistance to ZAP despite increasing its CpG content by 33% compared to its zoonotic ancestor, SIVsmm. This evasion of ZAP is mediated by changes in the 3’ U3 LTR region that promote HIV-2 fitness in primary human T cells. Our results show that ZAP contributes IFN mediated response against SIVsmm in human cells and show the versality of primate lentiviruses in evading ZAP-mediated antiviral immunity. Furthermore, our findings suggest that evolution of an efficient ZAP evasion strategy allows viruses to tolerate increased genomic CpG levels. Thus, our findings demonstrate that ZAP poses a barrier for zoonotic viruses, and that evolution of ZAP evasion strategies can lower the pressure driving CpG suppression in their genomes.

The link between viral CpG suppression and ZAP evasion is well established (7, 13, 50, 56–58, 60–62). However the determinants of ZAP sensitivity are complex and include not only the CpG number, but also their localization, context and local RNA structure (13, 60, 62). We have previously shown that CpG frequency in the 5’ end of the *env* gene determines the sensitivity of primary HIV-1 strains to ZAP (13). However, HIV-2 *env* plays no role in its resistance to ZAP, which underscores the divergent evolutionary paths taken by HIV-1 and HIV-2 in adapting to human immune defences and evading ZAP. In case of SIVsmm, ZAP efficiently decreased both Env and Gag levels, which are expressed from independent, spliced and unspliced viral mRNA, respectively. The broad viral transcript targeting by ZAP can be explained by the fact that all HIV-2 and SIVsmm RNA splicing products contain the untranslated 3’ U3 LTR region. Analyses of chimeric HIV-2/SIVsmm recombinants and iCLIP revealed that this RNA region is specifically targeted by ZAP. Furthermore, our findings suggest that while the *nef* ORF contributes to ZAP evasion due to its overlap with 3’ U3 LTR, the Nef protein does not act as a direct antagonist of ZAP. While the immunomodulatory functions of HIV-2 and SIVsmm Nefs are highly conserved (16, 63, 64), changes it its coding region can additionally promote viral adaptation to humans and resistance to antiviral restriction factors such as ZAP.

ZAP contributes to restriction of many animal-infecting RNA viruses (65–69) (66, 70, 71),. Here, we found that ZAP contributes to IFN-mediated restriction of SIVsmm in primary human T cells. Therefore, ZAP constitutes a part of the interspecies barrier protecting humans from zoonotic pathogens. ZAP targets the UTRs not only of SIVsmm but also several viral and host mRNAs (7, 72–74). It remains to be established if the UTR region is commonly recognized and targeted by ZAP in other viruses and whether pathogens with increased CpG content other than HIV-2 utilize an CpG-independent ZAP evasion strategy.

While our results identify ZAP as an effective inhibitor of SIVsmm replication in human T cells during IFN-response activation, it is not the only antiviral restriction factor that represents a cross-species barrier to this zoonotic virus. For example, it has been shown that SIVsmm is less efficient at counteracting human APOBEC3F (26), IFI16 (75) and GBP5 than HIV-2 (26, 75–77), suggesting that the latter had to evolve multiple innate immune evasion strategies to spread significantly in humans. Apart from ZAP, type I IFN upregulates the expression of hundreds of other ISGs (78–80), most of which have never been studied in the context of HIV-2 or SIVsmm restriction. Deciphering the contribution of all individual human ISGs to SIVsmm restriction would be an ambitious and time-consuming task that lies beyond the scope of the current study. Nevertheless, the generated chimeric HIV-2/SIVsmm infectious molecular clones will be highly useful for further studies investigating the innate immune mechanisms limiting SIVsmm cross-species transmission and spread in humans.

We found that the U3 region of HIV-2 accumulated changes during its human-specific adaptation that result in changes in the second half of the *nef* ORF, contribute to replicative fitness and mediate ZAP’s resistance of HIV-2. The corresponding 5’ part of the U3 region of SIVsmm was significantly bound by ZAP, and its introduction into HIV-2 7312A could confer partial ZAP sensitivity. It remains to be clarified how the structure of this RNA region affects the recognition by ZAP and how changes in this region affect the replication and ZAP sensitivity of other groups of HIV-2 and SIVs. Notably, the U3 LTR regions of HIV-2 isolates have extremely high genomic variability (81). Considering the presence of this region in both the unspliced RNA and all major splicing products, it is surprising that so little is known about the physiological role of the U3 region found upstream of the major Sp1 and NF-κB transcription factor binding sites (81). Our finding that this poorly-studied RNA region enables evasion of an IFN-induced antiviral restriction factor uncovers a new role of the HIV-2 U3.

SIVsmm replicates in human cells but is not well equipped to evade IFN-mediated response in humans. The contribution of ZAP to this restriction might at first seem counter-intuitive, as sooty mangabeys express a highly active form of ZAP, and SIVsmm has co-evolved with its host much longer than HIV-2. However, the difference in the antiviral immune response activation can may explain this phenomenon: sooty mangabeys infected by SIVsmm have high viral loads but do not develop the excessive immune activation and inflammation leading to a progressive CD4+ T cell loss and immunodeficiency (17, 39, 82–84). This is likely a result of a long pathogen-host co-evolution and the lack of fully functional TLR4 receptor in sooties (82). In contrast, HIV-2 infection in humans is characterized by high levels of immune activation and IFN production that upregulates the expression of ZAP and its cofactor TRIM25 (2, 9, 66, 68). Our results support a model in which epidemic HIV-2 had to evolve increased fitness and IFN resistance during its adaptation to humans. A part of this human-specific adaptation was mediated by mutations in the 3’ U3 LTR region that increase replicative fitness in human T cells and allow evasion of ZAP through a CpG-independent mechanism. This adaptation reduced the selective pressure acting against CpGs in the HIV-2 genome, leading to a substantial accumulation of these rare dinucleotide over the course of its evolution.

In conclusion, we show that ZAP contributes to the IFN-mediated restriction of SIVsmm in human T cells and in part elucidate how HIV-2 evolved to evade it. By demonstrating that HIV-2’s resistance to ZAP is mediated by a CpG-independent mechanism involving the 3’ LTR U3 region, we identified a novel ZAP evasion mechanism, that requires further molecular characterization. It remains to be clarified whether ZAP evasion contributes to the fitness of HIV-2 *in vivo*, as well as if it removes the evolutionary constraints posed by ZAP on CpG content in lentiviral genomes. It will be interesting to test if untranslated regions of other viruses are also targeted by ZAP and if other successful zoonotic viral pathogens evolved similar evasion mechanisms. Thus, further studies on the interplay between ZAP and viral RNAs will provide insights into the virus-host arms race and the mechanisms shaping viral evolution.

## ACKNOWLEDGEMENTS

We thank Regina Burger, Jana Romana Fischer, Daniela Krnavek, Martha Mayer, Birgit Ott, Kerstin Regensburger and Nicola Schrott for laboratory assistance. Prof. Paul Bieniasz and Prof. Molly Ohainle kindly provided the HEK293T ZAP KO and Jurkat ZAP KO cells. TZM-bl cells have been provided by J. C. Kappes and X. Wu and Tranzyme Inc. through the NIH AIDS Reagent Program. The p27 antibody provided by Dr. Karen Kent and Caroline Powell was obtained through the NIH HIV Reagent Program, Division of AIDS, NIAID, NIH and the Centre for AIDS Reagents, NIBSC, UK, supported by EURIPRED (EC FP7 INFRASTRUCTURES-2012 – INFRA-2012-1.1.5: Grant Number 31266). The HIV-2 Env antiserum provided by Dr. Raymond Sweet, SmithKline Beecham Pharmaceuticals was obtained through the same NIH program. We also thank Prof. Dennis Kätzel, Prof. Daniel Sauter and Prof. Konstantin Sparrer for critical reading of the article and helpful discussions.

## AUTHOR CONTRIBUTIONS

Dorota Kmiec: Conceptualization, Experimental work, Formal analysis, Methodology, Validation, Writing—original draft. Rayhane Nchioua: Experimental work, Formal analysis, Methodology. Alexander Gabel: Formal analysis, Methodology. Asimenia Vlachou: Experimental work, reagent generation. Sabina Ganskih: Experimental work. Sümeyye Erdemci-Evin: Reagent generation. Stacey Lab: Reagent generation. Diane Carnathan: Reagent generation. Steven Bosinger: Reagent generation. Ben Berkhout: Formal analysis. Atze T Das: Formal analysis, Visualization. Mathias Munschauer: Formal analysis, Visualization. Frank Kirchhoff: Conceptualization, Formal analysis. All authors: writing—review & editing.

## SUPPLEMENTARY DATA

Supplementary Data are available at NAR online.

## DATA AVAILABILITY

The data generated by iCLIP has been deposited to GEO repository under the identifier GSE290608 and will be released to the public upon publication of the manuscript. The primary data underlying this article as well as generated resources will be shared on a reasonable request to the corresponding author.

