## Supplementary Figures for "HIV-2 evades restriction by ZAP through adaptations in the U3 LTR region despite increased CpG levels"

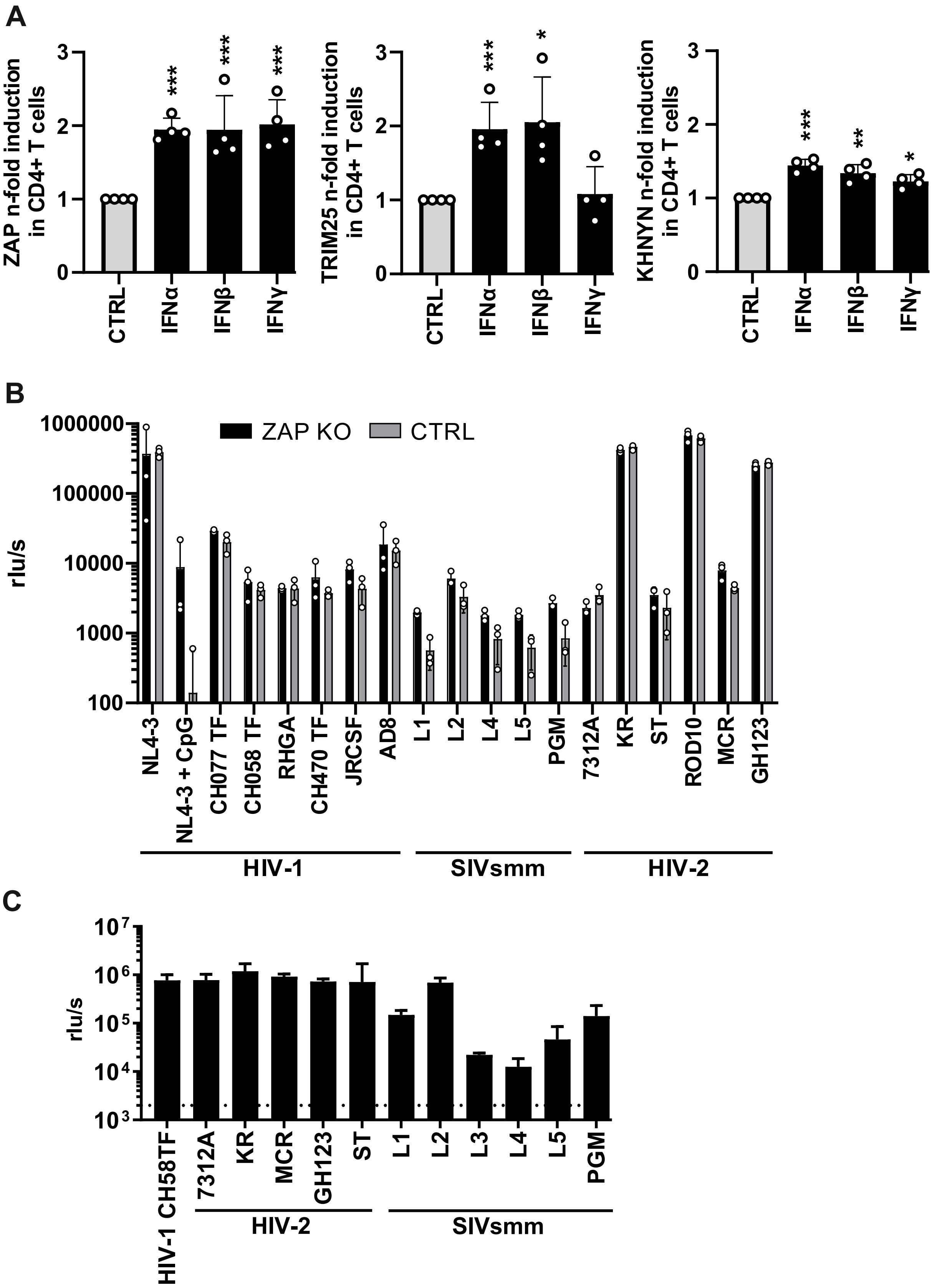
A
ZAP n-fold induction in CD4+ T cells
TRIM25 n-fold induction in CD4+ T cells
KHNYN n-fold induction in CD4+ T cells
CTRL
IFNα
IFNβ
IFNγ
B
ZAP KO
CTRL
rlu/s
NL4-3
NL4-3 + CpG
CH077 TF
CH058 TF
RHGA
CH470 TF
JRCSF
AD8
L1
L2
L4
L5
PGM
7312A
KR
ST
ROD10
MCR
GH123
HIV-1
SIVsmm
HIV-2
C
rlu/s
HIV-1 CH58TF
7312A
KR
MCR
GH123
ST
L1
L2
L3
L4
L5
PGM
HIV-2
SIVsmm

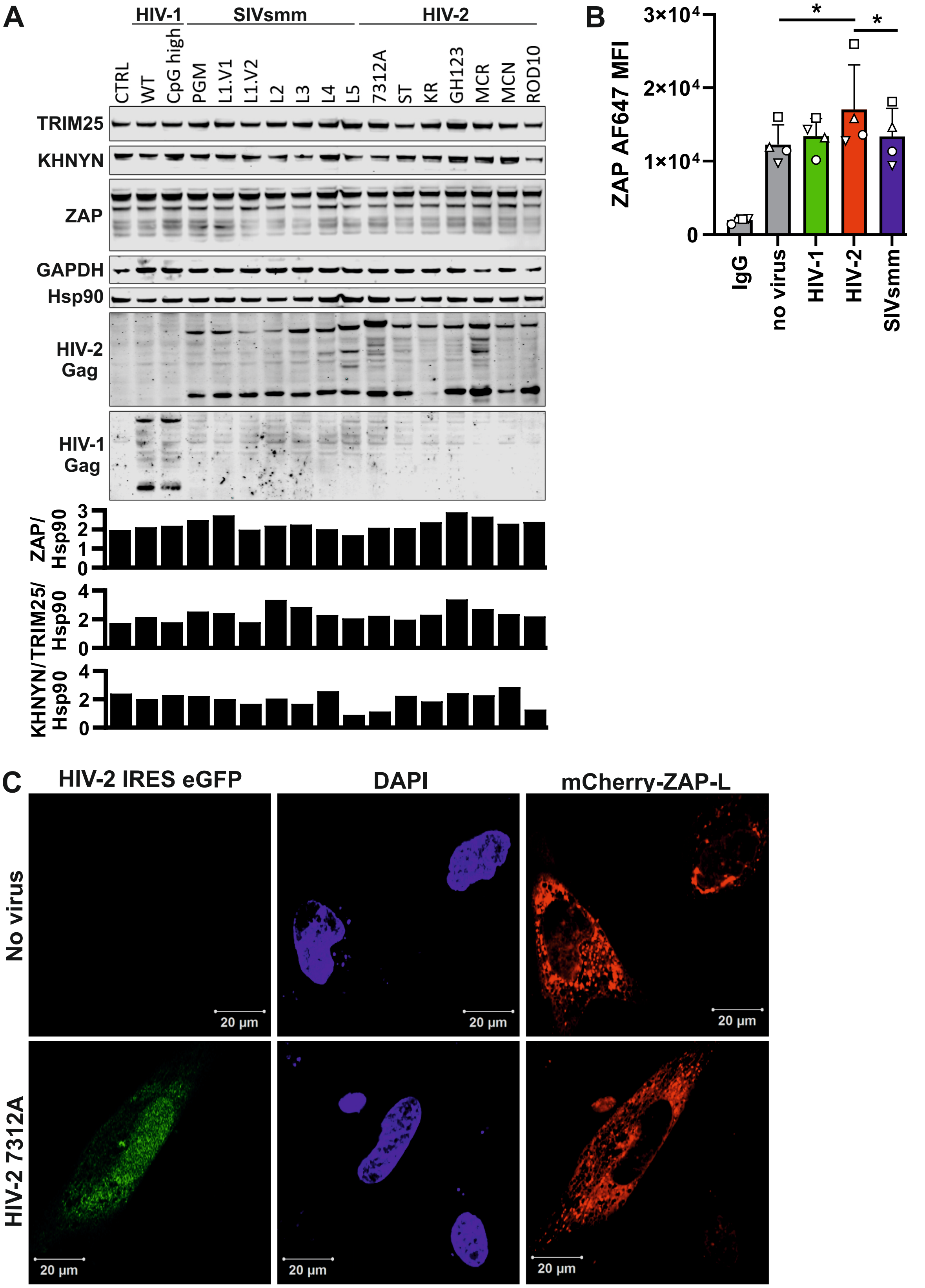
A
HIV-1
SIVsmm
HIV-2
CTRL
WT
CpG high
PGM
L1.V1
L1.V2
L2
L3
L4
L5
7312A
ST
KR
GH123
MCR
MCN
ROD10
TRIM25
KHNYN
ZAP
GAPDH
Hsp90
HIV-2 Gag
HIV-1 Gag
ZAP/ Hsp90
TRIM25/ Hsp90
KHNYN/ Hsp90
B
ZAP AF647 MFI
3×10⁴
2×10⁴
1×10⁴
0
*
*
IgG
no virus
HIV-1
HIV-2
SIVsmm
C
HIV-2 IRES eGFP
DAPI
mCherry-ZAP-L
No virus
HIV-2 7312A
20 µm

## A

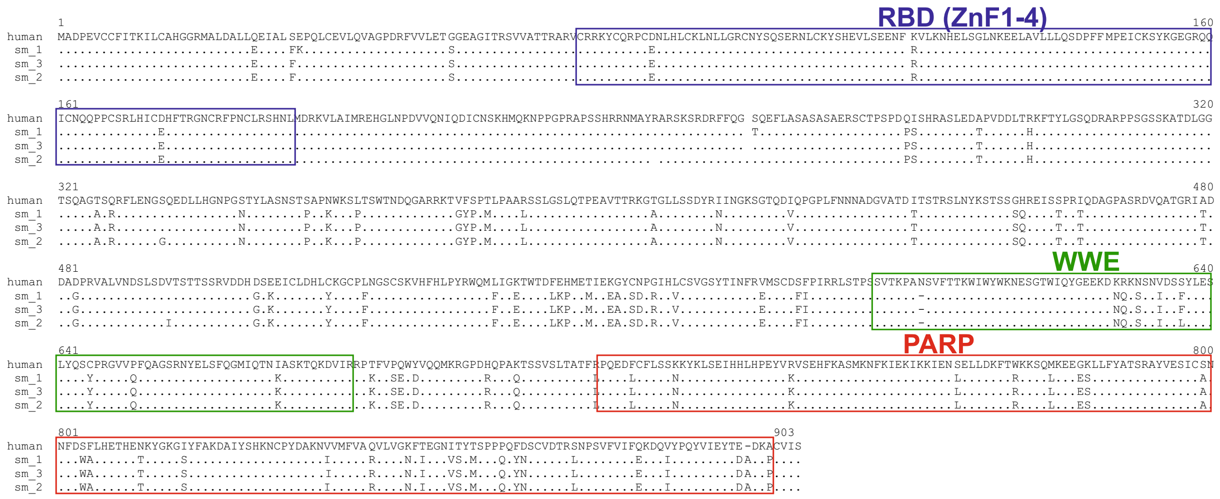
RBD (ZnF1-4)
WWE
PARP
human
sm_1
sm_3
sm_2
1
160
161
320
321
480
481
640
641
800
801
903

**Figure S4** Kmiec et al., 2025

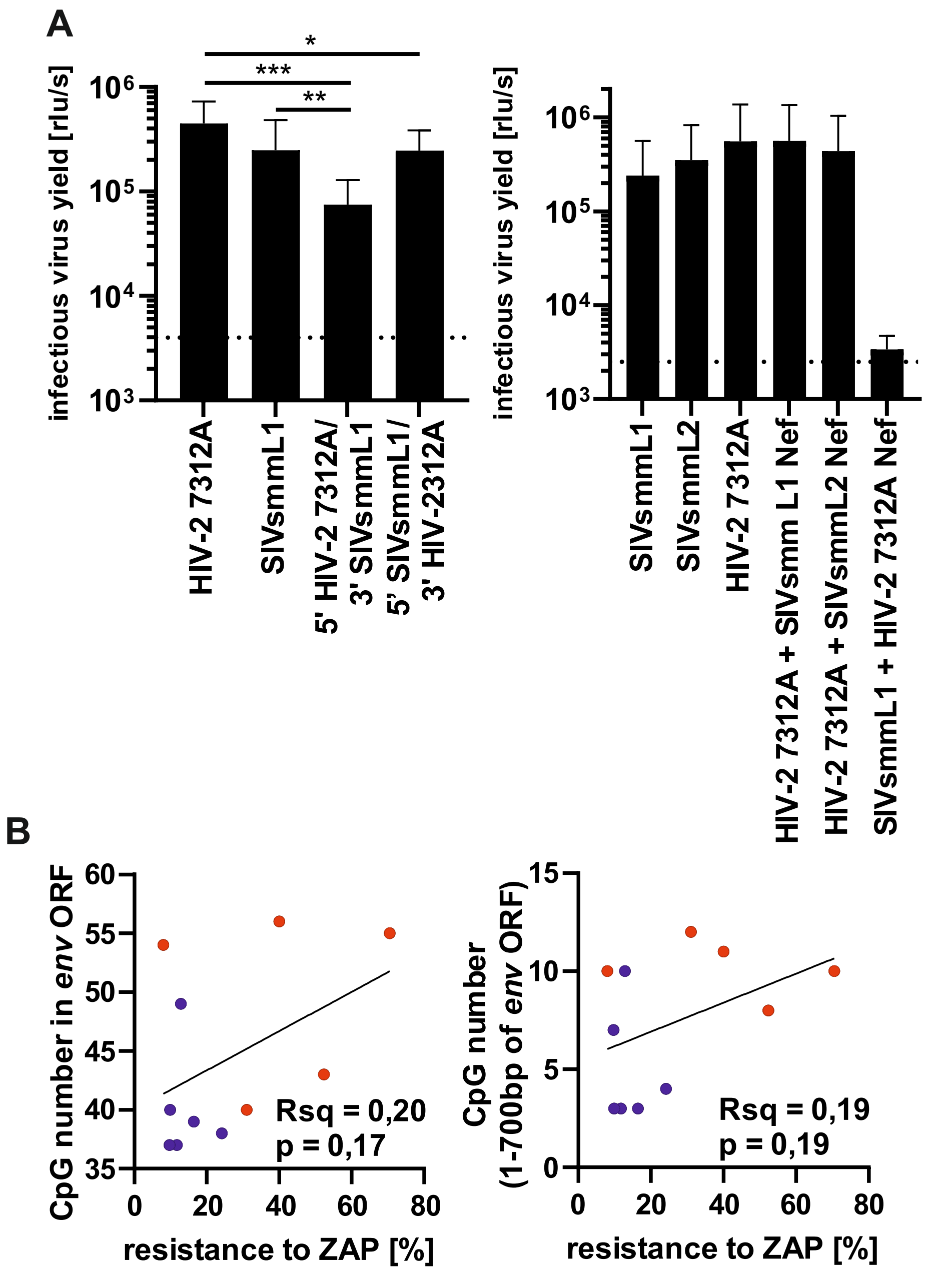

### Figure S5



```
        1                                                                                                   100
7312A   TGGAAGGGATTTTTTATAGTGAAAGAAGACATAGGATATTAGATACATATTTTGAGAATGAGAGAGGTATAGTTGGTGGATGGCAGAACTACACATATGG
  PGM   TGGAAGGGATTTATTACAGTGAGAGAAGACATAGAATCTTAGACATATACTTAGAAAAGGAAGAAGGCATCACCCCAGATTGGCAGAATTACACATCAGG
   L1   TGGAAGGGATTTATTACAGTGAAAGGAGACATAAAATATTAGATCTGTACTTAGAAAAGGAAGAAGGAATAGTACCAGATTGGCAAAATTACACATCAGG
   L2   TGGAAGGGATTTATTACAGTGATAGGAGACATAAAATATTAGATCTGTACTTAGAAAAGGAAGAAGGAATAGTACCAGATTGGCAAAATTACACATCAGG
   L3   TGGAAGGGATATATTACAGTGAAAGAAGGCATAAAATATTAGATTTATACCTAGAAAAAGAAGAAGGGATCATTCCTGATTGGCAGAATTACACAAAAGG
   L4   TGGAAGGGATTTATTACAGTGAGAGGAGACATAGAATATTAGACATATACCTAGAAAAGGAAGAAGGAATAATACCTGATTGGCAGAATTACACCTCAGG
   L5   TGGAAGGGATTTATTACAATCAGAGAAGACATAGAATACTAGACATTTACTTAGAAAAGGAGGAAGGAATCATCCCTGACTGGCAGAATTACACATCAGG

        101                                                                                                 200
7312A   GCCAGGGATAAGATACCCAAAACACTTTGGCTGGCTGTGGAAACTAGTGCCAGTGGAAGTGGCAGCAGTAACCCGGGAGGAAGAGGAGACCCATTGTCTA
  PGM   ACCAGGAATCAGATACCCAATGTTCTTTGGCTGGCTATGGAAATTGGTCCCAGTAAGTGCCTCAGATGAAGCTC---AAGAAGACGAGACACATTACTTA
   L1   GCCAGGAACCAGATATCCTAAGGTCTTTGGATGGCTCTGGAAATTAGTCCCAGTAAATGTCTCAGATGAAGCTC---AGGAAGATGAGACACATCGCTTG
   L2   GCCAGGAATCAGATATCCTAAGGTCTTTGGATGGCTCTGGAAATTAGTCCCAGTAAATGTCTCAGATGAAGCTC---AGGAAGATGAGACACATTGTTTG
   L3   GCCAGGGATAAGATACCCAATGACGTTTGGGTGGCTATGGAAGCTAGTGCCAGTAAATGTATCAGATGAGGCAG---AAACAGATGAGACCCACTGTCTG
   L4   GCCAGGAATAAGATATCCAATGTGCTTTGGATGGTTGTGGAAATTAGTACCAGTAGATGTCTCGGATGAGGCAC---AAGAAGATGAGACACACTATCTG
   L5   ACCAGGAATAAGATACCCAATGATGTTCGGATGGCTGTGGAAACTAGTCCCAGTAGAGGTCTTAGATGAGGCAC---AGAATGATGAGACACATTGTCTG

        201                                                                                                 300
7312A   GTGCACCCAGCACAGACCGCCGCATGGGATGATCCCCATGGGGAGACTCTTGTCTGGCAGTTTGATTCCCTCCTGGCATATTCAGATGAGGCCTTCAACA
  PGM   GTGCATCCAGCTCAGATATCCCAGTGGGATGATCCCTGGGGAGAGGTTCTGGCATGGAAGTTTGATTCACAACTAGCCTACAGATATGAGGCTTTCATTA
   L1   ATGCATCCAGCACAAACTTCTCAGTGGGATGACCCCTGGGGAGAGGTATTGGCATGGAAGTTTGATCCAGAGTTAGCTTATAACTACAAGGCATTTGTTA
   L2   ATGCATCCAGCACAAATTTCTCAGTGGGATGACCCCTGGGGAGAGGTATTGGCATGGAAGTTTGATCCAGAGTTAGCTTACAATTATAAGGCATTTATTG
   L3   GTACATCCGGCAGGGACAAGCAAGTGGGATGATCCGTGGGGTGAAACCCTAGCATGGAGATTTGATCCAACACTGGCTCATAGCTATGAAGCATTCATTA
   L4   GTGCATCCTGCACAGACAAGTCAGTGGAATGACCCATGGGGAGAAGTTCTAGCTTGGAAGTTTGATCCCACATTAGCTTATACTTATGAGGCATTTGTTA
   L5   GTGCATCCAGCACAGACGGGCCAATGGGATGACCAGTGGGGAGAGGTCTTGGCATGGAAGTTTGACCCCACTCTGGCTTATACCTATGAGGCATATATTA

                                                                                            nef STOP
        301                                                                                                 400
7312A   GGTTCCCAGAAGAGTTTGGGTATCAGTCAGGATTACCAGAGAAGGAATGGAAGGCTAGACTAAAAGCAAGAGGCATACCTACAGAGTAGACAGGAAAGCA
  PGM   GACACCCAGAAGAGTTTGGCAGTAAGTCAGGCTTGTCAGAGGAAGAGGTAAAGAGAAGGCTAACCGCAAGAGGCCT------------------------
   L1   AGTACCCAGAAGAGTTTGGTAGTATGTCAGGATTGCCAGAGGAAGAGGTAAAGAGAAGGCTAACCGCAAGAGGCCT------------------------
   L2   AGCACCCAGAAGAGTTTGGTAGTATGTCAGGCTTGTCAGAGGAAGAGGTGAAGAGAAGGCTAACCGCAAGAGGCCT------------------------
   L3   GATACCCAGAGGAGTTTGGATGGAGCTCAGGGCTGTCACAGGAAGAGGTTGAGAGAAGGCTAACCGCAAGAGGCCT------------------------
   L4   AATATCCAGAAGAATTTGGTAGCAAGTCAGGCTTGTCAGAGGAAGAGGTTAAGAGAAGGCTAACCGCAAGAGGCCT------------------------
   L5   AGTACCCAGAAGAGTTTGGGGACAAGTCAGGCTTGTCAGAGGAAGAGGTTAAGAGAAGGCTAACCGCAAGAGGCCT------------------------

                                                    nef STOP
        401                                                                                                 500
7312A   GCAGCATAAAGAGGAACTAGCTGACGCTGCATAAGAAAGGAAACTGGCTGACACTGCAGGGACTTTCCAGAAGGGGCTGTAACAGTGGGAGGGACATGGG
  PGM   ----TTTAAAAA-----TGGCTGAC--------AAGAAGGAAACAAGCTGAGACAGCAGGGACTTTCCACAAAGGG-GATGTTACGGGGAGGTACTGGGG
   L1   ----TTTAAAAA-----TGGCTAAC--------AAGAAGGAAACAAGCTGAGACAGCAGGGACTTTCCACAAAGGG-GATGTTATGGGGAGGTACTGGGG
   L2   ----TTTAAAAA-----TGCCTGAC--------AAGAAGGAAACAAGCTGAGACAGCAGGGACTTTCCACAAAGGG-GATGTTATGGGGAGGTACCGGGG
   L3   ----TTTAAAGA-----TGGCTGAC--------AAGGGAGGAAACAGCTGAGACAGCAGGGACTTTCCACAAGGGGCTGTAACA-GGGGAGGTACTTGGG
   L4   ----CTTAAAAA-----TGGCTGAC--------AAGAAGGAAACTAGCTGAGACAGCAGGGACTTTCCACAA-------------GGGGATGTTATGGGG
   L5   ----TTTAAAAA-----TGGCTGAC--------AAGAAGGAAACTAGCTGAGACTGCAGGGACTTTCCACAGTGGGAGGCATCAATGGGAGGTAACTGGG

        501                                                      558
7312A   AGGAGTTGGTGGGGAACGCCCATTAAGCCTCTGTATAAATGTACCCGCTTCTTGCATT
  PGM   AGGAGCTGGCTGG-AACGCCCACTTATTTTCTGTATAAATACAACTGCATTTCGCTCT
   L1   AGGAGCTGGCTGG-AACGCCCACTTATTCTCTGTATAAATACAACTGCATTTCGCTCT
   L2   AGGAGCTGGCTGG-AACGCCCACTTATTCTCTGTATAAATACAACTGCATTTCGCTCT
   L3   AGGAGCTGGTGGGGAACGCCCA-TTATTCTCTGTATAAATGTAGCTGCTGAACGCTCT
   L4   AGGAGCTGGTCGGGAACGCCCACTTTCTTGATGTATAAATACAACTGCATTTCGCTCT
   L5   AGGTACTGGTGGGGAACGCCCATTTATTCTCTGTATAAATACAACCGCTTGTCGCTCT
```

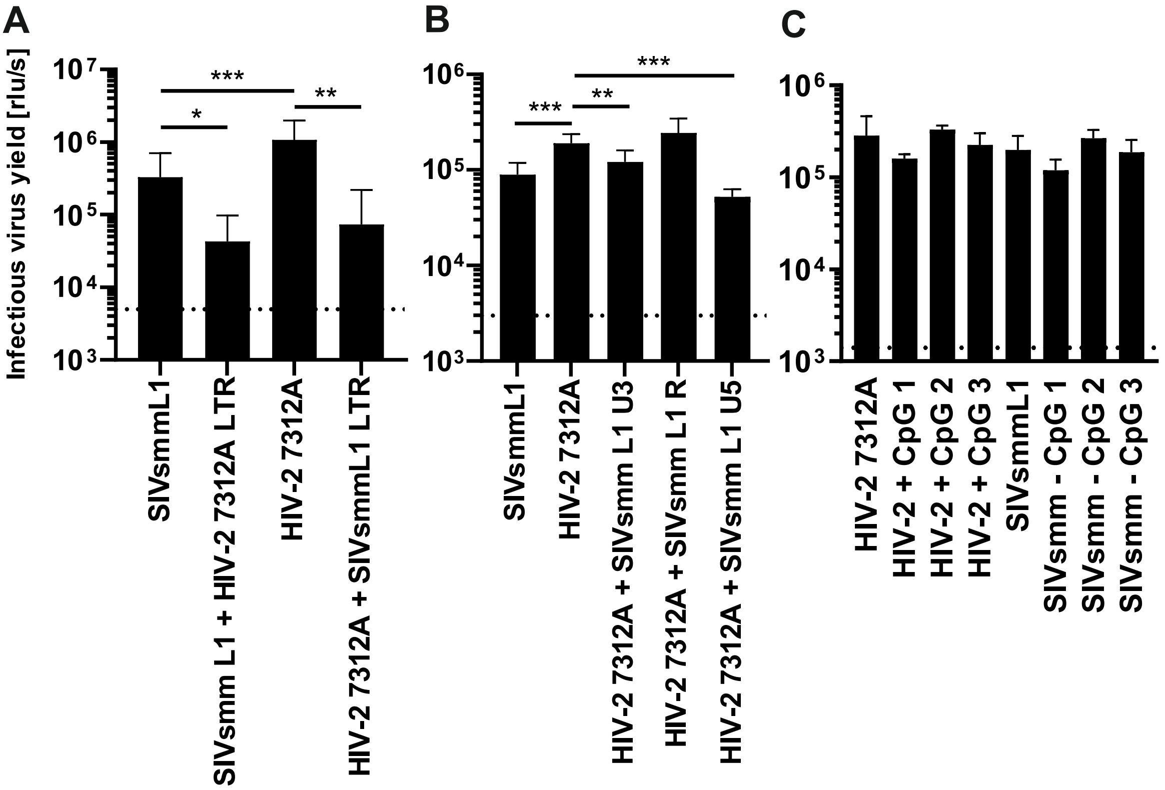
A
B
C
Infectious virus yield [rlu/s]
10^7
10^6
10^5
10^4
10^3
***
*
**
***
**
SIVsmmL1
SIVsmm L1 + HIV-2 7312A LTR
HIV-2 7312A
HIV-2 7312A + SIVsmmL1 LTR
HIV-2 7312A + SIVsmm L1 U3
HIV-2 7312A + SIVsmm L1 R
HIV-2 7312A + SIVsmm L1 U5
HIV-2 + CpG 1
HIV-2 + CpG 2
HIV-2 + CpG 3
SIVsmm - CpG 1
SIVsmm - CpG 2
SIVsmm - CpG 3

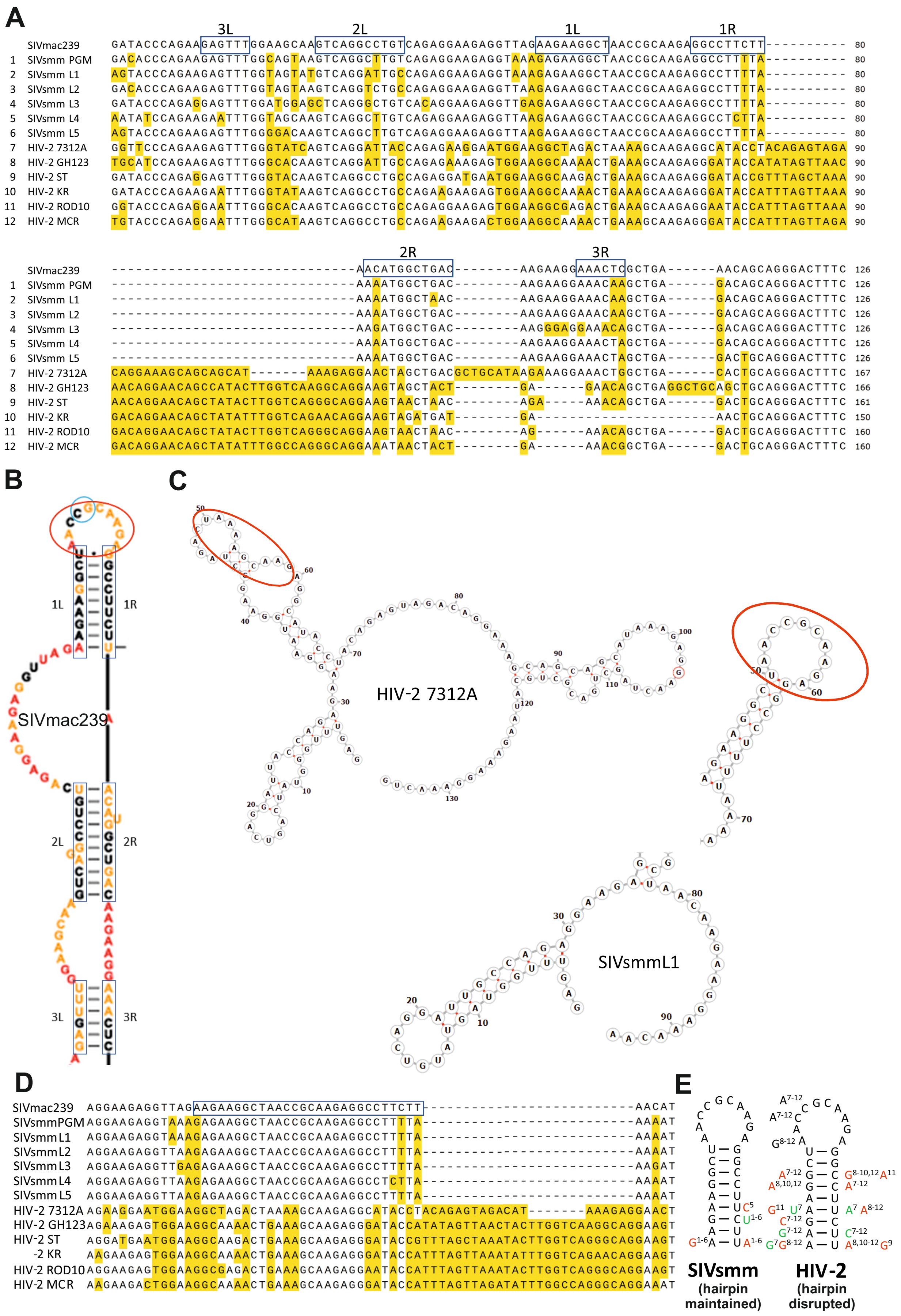
A
3L 2L 1L 1R
SIVmac239 GATACCCAGAAGAGTTTGGAAGCAAGTCAGGCCTGTCAGAGGAAGAGGTTAGAAGAAGGCTAACCGCAAGAGGCCTTCTT---------- 80
1 SIVsmm PGM GACACCCAGAAGAGTTTGGCAGTAAGTCAGGCCTTGTCAGAGGAAGAGGTAAAGAGAAGGCTAACCGCAAGAGGCCTTTTA---------- 80
2 SIVsmm L1 AGTACCCAGAAGAGTTTGGTAGTATGTCAGGATTGCCAGAGGAAGAGGTAAAGAGAAGGCTAACCGCAAGAGGCCTTTTA---------- 80
3 SIVsmm L2 GACACCCAGAAGAGTTTGGTAGTAAGTCAGGTCTGCCAGAGGAAGAGGTTAAGAGAAGGCTAACCGCAAGAGGCCTTTTA---------- 80
4 SIVsmm L3 GATACCCAGAGGAGTTTGGATGGAGCTCAGGGCTGTCACAGGAAGAGGTTGAGAGAAGGCTAACCGCAAGAGGCCTTTTA---------- 80
5 SIVsmm L4 AATATCCAGAAGAATTTGGTAGCAAGTCAGGCTTGTCAGAGGAAGAGGTTAAGAGAAGGCTAACCGCAAGAGGCCTCTTA---------- 80
6 SIVsmm L5 AGTACCCAGAAGAGTTTGGGGACAAGTCAGGCTTGTCAGAGGAAGAGGTTAAGAGAAGGCTAACCGCAAGAGGCCTTTTA---------- 80
7 HIV-2 7312A GGTTCCCAGAAGAGTTTGGGTATCAGTCAGGATTACCAGAGAAGGAATGGAAGGCTAGACTAAAAGCAAGAGGCATACCTACAGAGTAGA 90
8 HIV-2 GH123 TGCATCCAGAAGAGTTTGGGCACAAGTCAGGATTGCCAGAGAAAGAGTGGAAGGCAAAACTGAAAGCAAGAGGGATACCATATAGTTAAC 90
9 HIV-2 ST GATACCCAGAGGAGTTTGGGTACAAGTCAGGCCTGCCAGAGGATGAATGGAAGGCAAGACTGAAAGCAAGAGGGATACCGTTTAGCTAAA 90
10 HIV-2 KR GATACCCAGAAGAATTTGGGTATAAGTCAGGCCTGCCAGAAGAAGAGTGGAAGGCAAAACTGAAAGCAAGAGGGATACCATTTAGTTAAA 90
11 HIV-2 ROD10 GGTACCCAGAGGAATTTGGGCACAAGTCAGGCCTGCCAGAGGAAGAGTGGAAGGCGAGACTGAAAGCAAGAGGAATACCATTTAGTTAAA 90
12 HIV-2 MCR TGTACCCAGAGGAATTTGGGCATAAGTCAGGCCTGCCAGAAGAAGACTGGAAGGCAAAACTGAAAGCAAGAGGGATACCATTTAGTTAGA 90
2R 3R
SIVmac239 ------------------------------AACATGGCTGAC--------AAGAAGGAAACTCGCTGA------AACAGCAGGGACTTTC 126
1 SIVsmm PGM ------------------------------AAAATGGCTGAC--------AAGAAGGAAACAAGCTGA------GACAGCAGGGACTTTC 126
2 SIVsmm L1 ------------------------------AAAATGGCTAAC--------AAGAAGGAAACAAGCTGA------GACAGCAGGGACTTTC 126
3 SIVsmm L2 ------------------------------AAAATGGCTGAC--------AAGAAGGAAACAAGCTGA------GACAGCAGGGACTTTC 126
4 SIVsmm L3 ------------------------------AAGATGGCTGAC--------AAGGGAGGAAACAGCTGA------GACAGCAGGGACTTTC 126
5 SIVsmm L4 ------------------------------AAAATGGCTGAC--------AAGAAGGAAACTAGCTGA------GACAGCAGGGACTTTC 126
6 SIVsmm L5 ------------------------------AAAATGGCTGAC--------AAGAAGGAAACTAGCTGA------GACTGCAGGGACTTTC 126
7 HIV-2 7312A CAGGAAAGCAGCAGCAT-------AAAGAGGAACTAGCTGACGCTGCATAAGAAAGGAAACTGGCTGA------CACTGCAGGGACTTTC 167
8 HIV-2 GH123 AACAGGAACAGCCATACTTGGTCAAGGCAGGAAGTAGCTACT--------GA------GAACAGCTGAGGCTGCAGCTGCAGGGACTTTC 166
9 HIV-2 ST AACAGGAACAGCTATACTTGGTCAGGGCAGGAAGTAACTAAC--------AGA-----AAACAGCTGA------GACTGCAGGGACTTTC 161
10 HIV-2 KR GACAGGAACAGCTATATTTGGTCAGAACAGGAAGTAGATGAT--------GA----------------------AACTGCAGGGACTTTC 150
11 HIV-2 ROD10 GACAGGAACAGCTATACTTGGTCAGGGCAGGAAGTAACTAAC--------AG------AAACAGCTGA------GACTGCAGGGACTTTC 160
12 HIV-2 MCR GACAGGAACAGCTATATTTGGCCAGGGCAGGAAATAACTACT--------GA------AAACGGCTGA------GACTGCAGGGACTTTC 160
B
1L 1R
SIVmac239
2L 2R
3L 3R
C
HIV-2 7312A
SIVsmmL1
D
SIVmac239 AGGAAGAGGTTAGAAGAAGGCTAACCGCAAGAGGCCTTCTT--------------------------AACAT
SIVsmmPGM AGGAAGAGGTAAAGAGAAGGCTAACCGCAAGAGGCCTTTTA--------------------------AAAAT
SIVsmmL1 AGGAAGAGGTAAAGAGAAGGCTAACCGCAAGAGGCCTTTTA--------------------------AAAAT
SIVsmmL2 AGGAAGAGGTTAAGAGAAGGCTAACCGCAAGAGGCCTTTTA--------------------------AAAAT
SIVsmmL3 AGGAAGAGGTTGAGAGAAGGCTAACCGCAAGAGGCCTTTTA--------------------------AAGAT
SIVsmmL4 AGGAAGAGGTTAAGAGAAGGCTAACCGCAAGAGGCCTCTTA--------------------------AAAAT
SIVsmmL5 AGGAAGAGGTTAAGAGAAGGCTAACCGCAAGAGGCCTTTTA--------------------------AAAAT
HIV-2 7312A AGAAGGAATGGAAGGCTAGACTAAAAGCAAGAGGCATACCTACAGAGTAGACAT-------AAAGAGGAACT
HIV-2 GH123 AGAAAGAGTGGAAGGCAAAACTGAAAGCAAGAGGGATACCATATAGTTAACTACTTGGTCAAGGCAGGAAGT
HIV-2 ST AGGATGAATGGAAGGCAAGACTGAAAGCAAGAGGGATACCGTTTAGCTAAATACTTGGTCAGGGCAGGAAGT
-2 KR AAGAAGAGTGGAAGGCAAAACTGAAAGCAAGAGGGATACCATTTAGTTAAATATTTGGTCAGAACAGGAAGT
HIV-2 ROD10 AGGAAGAGTGGAAGGCGAGACTGAAAGCAAGAGGAATACCATTTAGTTAAATACTTGGTCAGGGCAGGAAGT
HIV-2 MCR AAGAAGACTGGAAGGCAAAACTGAAAGCAAGAGGGATACCATTTAGTTAGATATTTGGCCAGGGCAGGAAAT
E
SIVsmm
(hairpin maintained)
HIV-2
(hairpin disrupted)

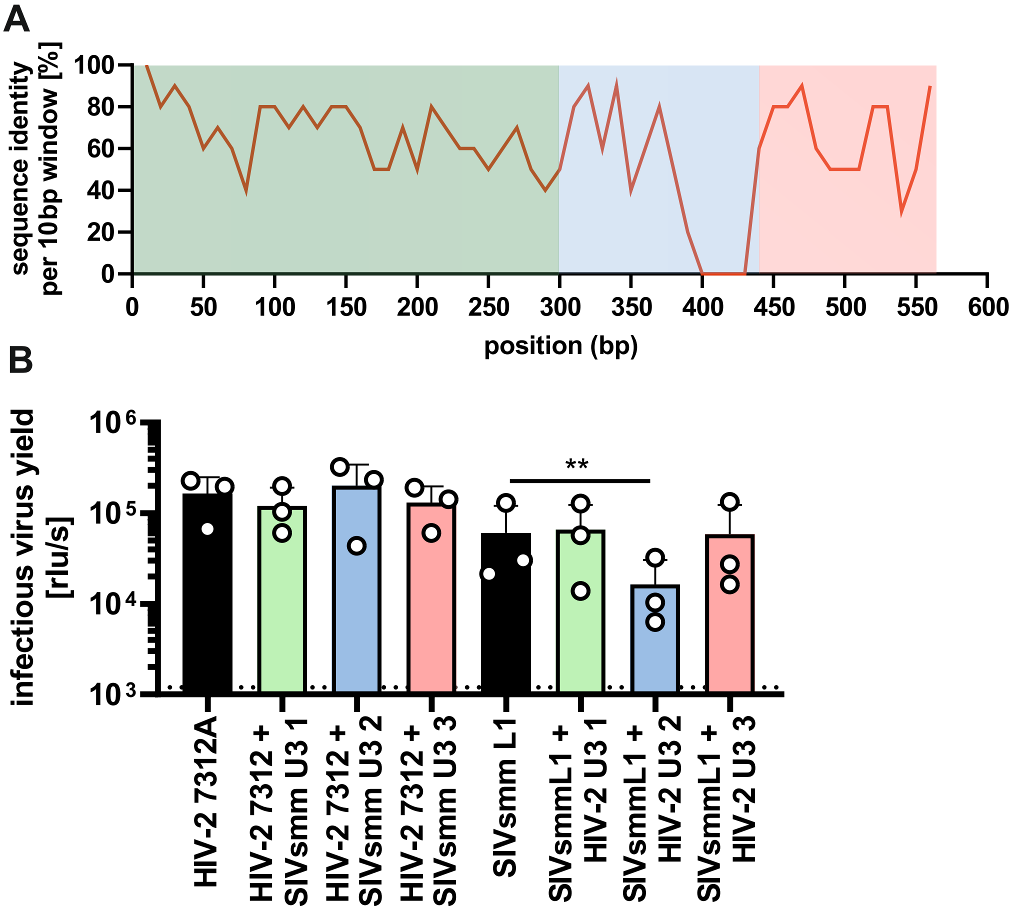
A
sequence identity per 10bp window [%]
100
80
60
40
20
0
0
50
100
150
200
250
300
350
400
450
500
550
600
position (bp)
B
infectious virus yield [rlu/s]
10^6
10^5
10^4
10^3
**
HIV-2 7312A
HIV-2 7312 + SIVsmm U3 1
HIV-2 7312 + SIVsmm U3 2
HIV-2 7312 + SIVsmm U3 3
SIVsmm L1
SIVsmmL1 + HIV-2 U3 1
SIVsmmL1 + HIV-2 U3 2
SIVsmmL1 + HIV-2 U3 3
