## Supplementary Figure legends for "HIV-2 evades restriction by ZAP through adaptations in the U3 LTR region despite increased CpG levels"

**Figure S1.** (A) Related to Figure 1 and 2. *Expression levels of ZAP and its cofactors TRIM25 and KHNYN in primary human PBMCs 2 days after stimulation with 500u/ml type I IFN (α, β) or 200u/ml type II IFN(γ) measured by flow cytometry. Each dot represents one donor. (B) Related to Figure 2. Raw infectivity values of HIV-1, HIV-2 and SIVsmm in transduced Jurkat CCRt cells and (C) transfected HEK293T ZAP KO cells.*

**Figure S2.** *Related to Figure 3.* ***(A)*** *Western blot of HEK293T cells transfected with indicated HIV-1, SIVsmm or HIV-2 provirus, and quantification of endogenous expression levels of ZAP, TRIM25 and KHNYN normalized to the Hsp90 housekeeping gene.* ***(B)*** *Expression levels of ZAP in pre-activated human PBMCs isolated from 4 donors, transduced with HIV-1, HIV-2 or SIVsmm and stained 3 days later with AF649-conjugated anti-ZAP and FITC-conjugated anti-p24 (Gag) antibodies. *, $p < 0.05$ calculated using paired Student's t-test.* ***(C)*** *Confocal microscopy images of HeLa cells co-transfected with HIV-2 IRES eGFP and mCherry-tagged ZAP and stained with nuclear stain (DAPI).*

**Figure S3.** *Related to Figure 3 and 4. (A) Protein alignment of human and sooty mangabey ZAP variants cloned from PBMC cDNA. RNA-binding domain containing 4 zinc-fingers (ZnF), WWE domain and catalytically inactive PARP domain are shown. Dots indicate amino acid conservation.*

**Figure S4.** *Related to Figure 4-6.* ***(A)*** *Raw infectivity values of chimeric HIV-2/SIVsmm half mutants and nef exchange mutant proviruses transfected into HEK293T ZAP KO cells. N=3 +SD.* ***(B)*** *Lack of significant correlation between CpG env number and ZAP resistance of HIV-2 (blue) and SIVsmm (red).*

**Figure S5.** *Related to Figure 6. Nucleotide alignment of U3 LTR regions of HIV-2 7312A and SIVsmmL1-L5. CpG dinucleotides are highlighted in green and the end of nef ORF is indicated by a black box.*

**Figure S6.** *Related to Figure 6.* ***(A)*** *Raw infectivity values of chimeric HIV-2/SIVsmm LTR mutant proviruses and 3'LTR U3, R and* ***(B)*** *U5 exchange mutants and* ***(C)*** *U3 CpG mutants in transfected into HEK293T ZAP KO cells.* N=3-5 + SD; *, $p < 0.05$; **, $p < 0.01$; ***, $p < 0.001$ *calculated using Student's t-test.*

**Figure S7.** Related to Figure 7. **(A)** An alignment of LTR U3 fragments of tested SIVsmm and HIV-2 proviruses. **(B)** Stem-loop structure in the SIVmac239 RNA based on SHAPE RNA structure probing. **(C)** RNA fold structure analysis of the 3L-3R sequence predicts a similar 1L-1R hairpin for SIVsmm L1 variant but not for HIV-2 7312A (but no 2L-2R or 3L-3R stem). CpG indicated by blue circle. Loop sequence indicated by red circle. **(D)** *Sequence alignment of the predicted loop region in the tested SIVsmm (labelled 1-6) and HIV-2 variants (7-12) with the SIVmac239 sequence. Non-conserved positions are highlighted in yellow. The blue box indicates the nucleotides that can fold a hairpin structure in the SIVmac239 RNA (based on SHAPE structure probing* (54). ***(E)*** *Sequence variation in the hairpin region in tested SIVsmm (left) and HIV-2 strains (right). The SIVmac239 RNA hairpin structure is shown with the nucleotides differing in the SIVsmm and HIV-2 strains indicated. The numbers in superscript refer to the labelled sequences shown in the bracket of panel D. Nucleotides in green allow basepairing, whereas nucleotides in red do not.*

**Figure S8.** *Related to Figure 8. (A) Sliding window (10bp) analysis of HIV-2 7312A and SIVsmmL1 nucleotide sequence identity based on aligned U3 sequences. Colors refer to U3 region A (green), B (blue) and C (pink). (B) Infectious virus yield of the WT and U3 mutants viruses in transfected HEK293T ZAP KO cells. N=3 + SD. **, P < 0.01; calculated using Student's t-test.*
